# Control and regulation of acetate overflow in *Escherichia coli*

**DOI:** 10.1101/2020.08.18.255356

**Authors:** Pierre Millard, Brice Enjalbert, Sandrine Uttenweiler-Joseph, Jean-Charles Portais, Fabien Létisse

## Abstract

Overflow metabolism refers to the production of seemingly wasteful by-products by cells during growth on glucose even when oxygen is abundant. Two theories have been proposed to explain acetate overflow in *Escherichia coli* – global control of the central metabolism and local control of the acetate pathway – but neither accounts for all observations. Here, we develop a kinetic model of *E. coli* metabolism that quantitatively accounts for observed behaviors and successfully predicts the response of *E. coli* to new perturbations. We reconcile these theories and clarify the origin, control and regulation of the acetate flux. We also find that, in turns, acetate regulates glucose metabolism by coordinating the expression of glycolytic and TCA genes. Acetate should not be considered a wasteful end-product since it is also a co-substrate and a global regulator of glucose metabolism in *E. coli*. This has broad implications for our understanding of overflow metabolism.

## Introduction

Overflow metabolism refers to the production of seemingly wasteful by-products by cells during growth on glycolytic substrates such as glucose, even when oxygen is abundant. Overflow metabolism has been reported for most (micro)organisms: yeasts produce ethanol (a phenomenon known as the Crabtree effect [1]), mammalian cells produce lactate (the Warburg effect in cancer cells [2]), and bacteria produce small organic acids such as acetate [3]. This ubiquitous phenomenon has been extensively investigated because of its fundamental and applied importance, but its origin and regulation remain to be clarified.

Acetate production by *Escherichia coli* has been studied for decades as a model of overflow metabolism. The recent rise of systems biology, which combines experimental techniques and mathematical modelling to achieve a mechanistic understanding of processes underlying the function of biological systems [4, 5], has provided a quantitative understanding of the cause of acetate overflow in *E. coli*. Acetate production is considered to result from an imbalance between *E. coli*’s capacities to produce and assimilate acetyl-CoA, the main precursor of acetate. Stoichiometric models suggest that this imbalance could be caused by various cell-level constraints, such as energy conservation [6, 7], recycling of cofactors [8, 9], membrane occupancy [10, 11], and resource allocation [12, 13]. These models – and the underlying theories – capture the essence of the overflow process and successfully predict the growth rate dependence of acetate production in *E. coli* [12–14].

However, none of these theories explain why increases in acetate concentration can interrupt or even reverse the acetate flux in *E. coli* [15], which is able to co-consume glucose and acetate even when glucose is abundant. Cell growth is not affected when acetate production is abolished [15], suggesting that this overflow is neither necessary for *E. coli* to sustain fast growth nor strictly imposed by the proposed constraints. This phenomenon cannot be reproduced by stoichiometric models since they do not account for metabolite concentrations. Kinetic models are therefore a necessary step towards a comprehensive understanding of the dynamics, control and regulation of metabolic systems. For example, a kinetic model of the Pta-AckA pathway has successfully been used to predict the reversal of the acetate flux at high acetate concentrations [15], a significant advance compared to stoichiometric models of acetate overflow. This effect is caused by thermodynamic control of the acetate pathway by the concentration of acetate itself, a mechanism that operates independently of enzyme expression. Still, thermodynamic control does not imply that enzymes exert no control whatsoever over acetate flux, though the degree of this control remains to be clarified. Moreover, this local mechanism does not explain the toxic effect of acetate on microbial growth [16, 17].

In this work, we used a systems biology approach to clarify the origin, control and regulation of acetate overflow in *E. coli* over the broad range of acetate concentrations this bacterium experiences under laboratory and industrial conditions as well as in its environmental niche [8, 18–25].

## Results

### Construction of a kinetic model of E. coli metabolism

Following a systems biology approach [5], we constructed a coarse-grained kinetic model of *E. coli* metabolism that links acetate metabolism with glucose uptake and growth (Figure 1A and Supplementary file 1). This model includes three processes: i) glucose transport and its conversion into acetyl-CoA by the glycolytic pathway, ii) utilization of acetyl-CoA in the TCA cycle (energy conservation) and anabolic pathways (production of building blocks) for growth, and iii) acetate metabolism, i.e. the reversible conversion of acetyl-CoA into acetate via phosphotransacetylase (Pta), acetate kinase (AckA), and an acetate exchange reaction between the cell and its environment.

**Figure 1.**
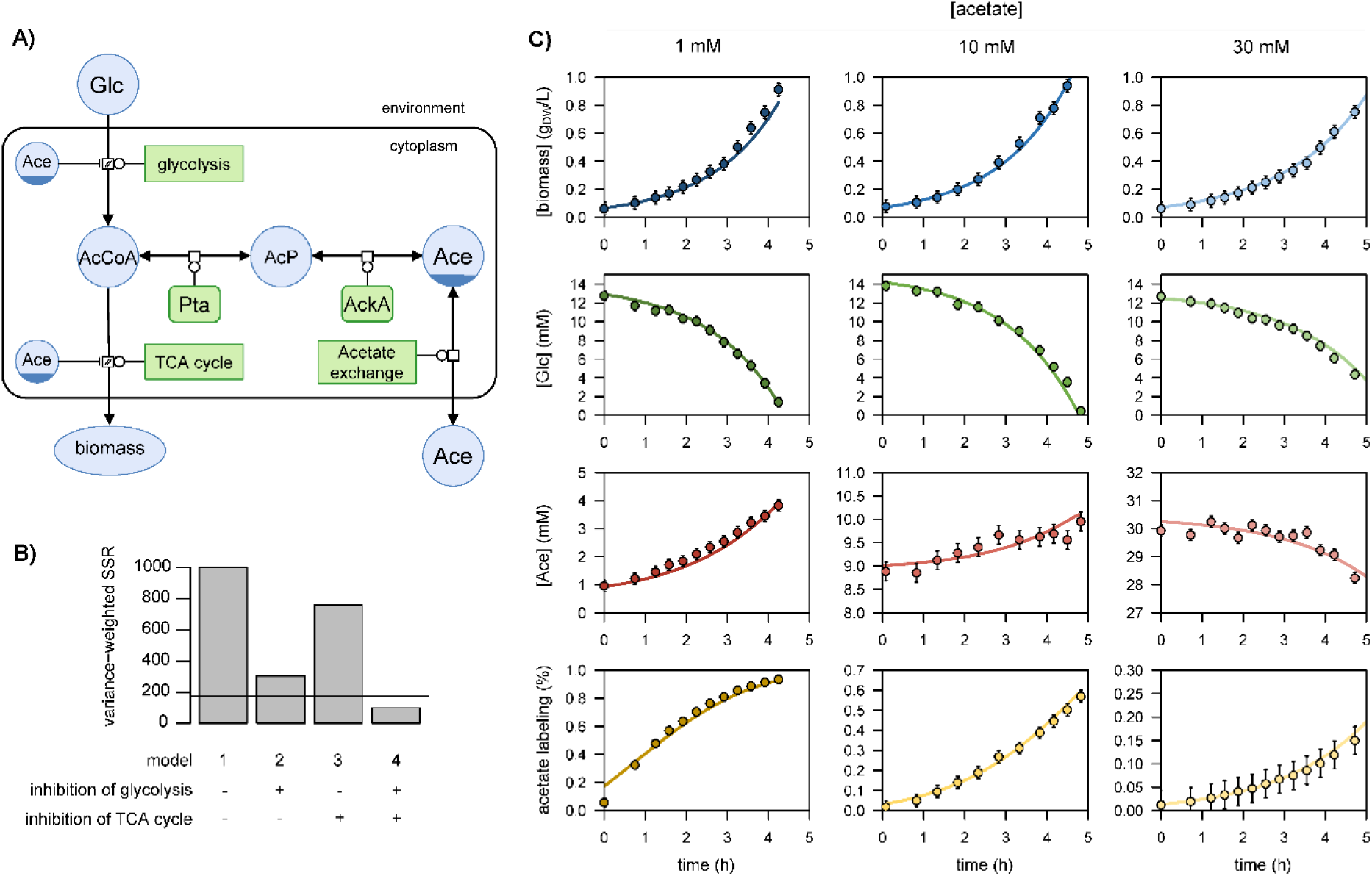
Representation of glucose and acetate metabolism in *Escherichia coli* (panel A), in Systems Biology Graphical Notation format (http://sbgn.org) [30]. We performed ^13^C-labeling experiments to calibrate the model and evaluated the goodness-of-fit for different topologies (panel B). The initial model (model 1), which does not include inhibition of the glycolytic pathway and TCA cycle by acetate, did not fit the data satisfactorily. Adding inhibition by acetate of glycolysis (model 2) or of the TCA cycle (model 3) improved the fit, but both pathways had to be inhibited (model 4) for the goodness-of-fit criterion to be satisfied. In panel B, the horizontal line represents the 95 % confidence threshold for the variance-weighted sum of squared residuals (SSR). The best fits of the experimental data obtained with model 4 are shown in panel C.

This initial model included 2 compartments, 6 species, 6 reactions and 29 parameters. The value of 19 parameters were taken directly from the literature [15, 26, 27] (Supplementary file 1). To estimate the remaining parameters, we performed three growth experiments on ^13^C-glucose (15 mM) plus different concentrations of ^12^C-acetate (1, 10 and 30 mM), as detailed in ref. [15]. These experiments provide information on the forward and reverse fluxes of acetate between the cell and its environment [15], and thereby improve the calibration of the model. The model was extended with isotopic equations [15, 28], and parameters were estimated by fitting concentration time courses of glucose, biomass, and acetate and of the ^13^C-enrichment of acetate (see *Materials and methods*). This initial model (model 1) did not fit the data satisfactorily (Figure 1B, *Materials and methods*). Adding inhibition of the glycolytic pathway by acetate (model 2) to account for the reduction in glucose uptake at high acetate concentrations improved the fit of the glucose concentration profile. Similarly, adding inhibition of the TCA cycle by acetate (model 3) improved the fit of the growth and acetate labelling profiles. A sufficiently accurate fit was only achieved however when both pathways were inhibited (model 4; Figure 1B, C). These modifications provide a mechanistic rationale for the reported inhibition of *E. coli* metabolism by acetate [15–17, 29].

### Acetate regulates glucose uptake, glycolysis, and the TCA cycle at the transcriptional level

A key hypothesis required for the model to fit the experimental data is that acetate inhibits the flux capacity of both the glycolytic pathway and the TCA cycle in *E. coli*. To determine whether this actually occurs *in vivo* at the transcriptional level, we monitored gene expression in *E. coli* grown on glucose (15 mM) plus acetate at different concentrations (0, 10, 50 or 100 mM). Transcriptomic results revealed a global and progressive remodelling of gene expression at increasing acetate concentrations (Figure 2A-B). We noted that the presence of acetate modulates the expression of genes involved in various biological functions: motility, biofilm formation, translation, stress response, metabolism, and transport of carbohydrates, amino acids and ions (Figure 2C). Importantly, the observed changes in gene expression cannot result solely from the inhibition of growth by acetate, as shown by comparing gene expression levels on glucose plus 100 mM acetate with those on glucose alone at the same growth rate (0.35 h^−1^) (Figure 2B).

**Figure 2.**
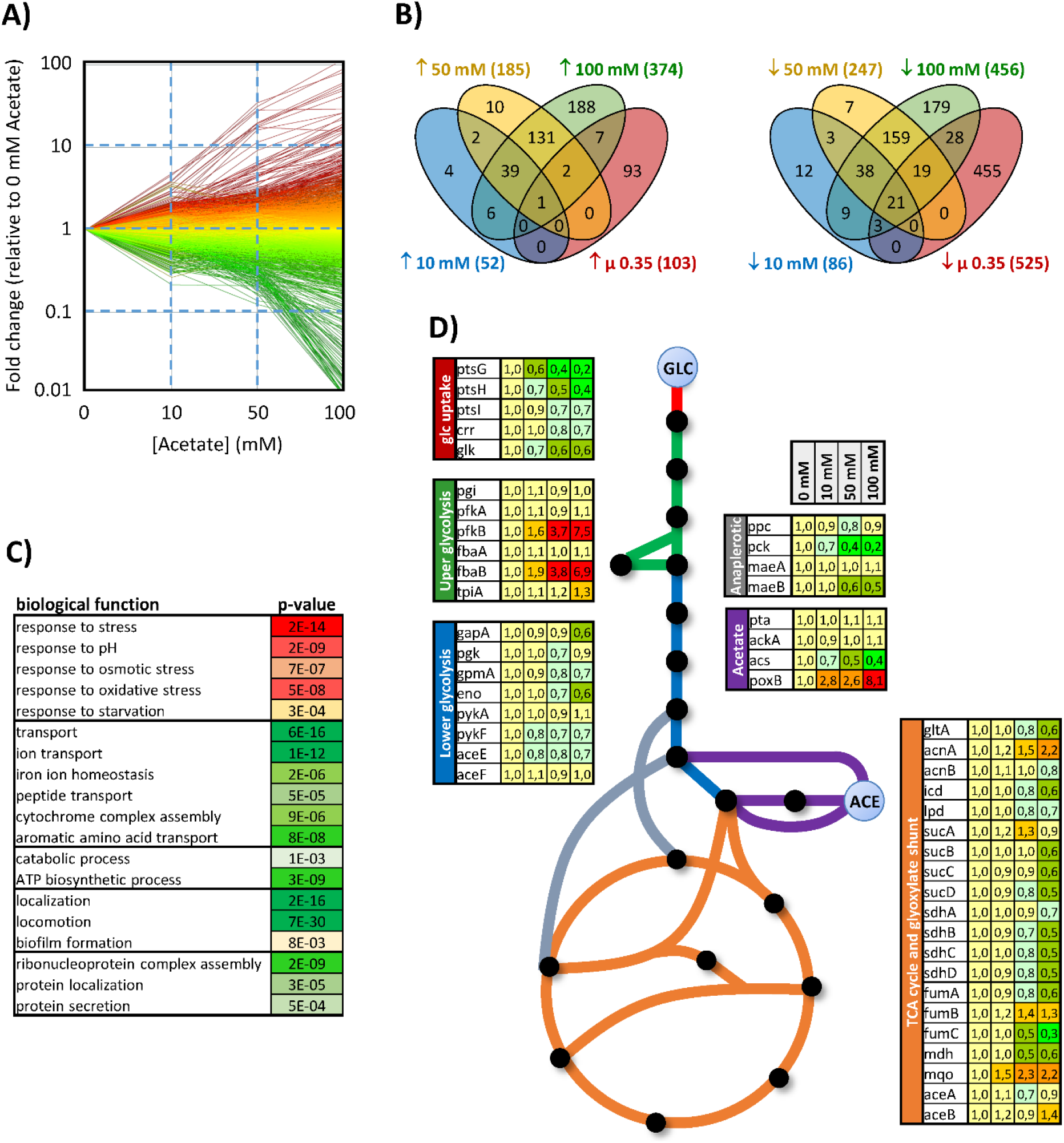
Response of the *E. coli* transcriptome to changes in acetate concentration (0, 10, 50 or 100 mM) during growth on glucose (15 mM). The changes in gene expression are shown in panel A. Each line represents the expression of a single gene relative to its expression level measured in the absence of acetate. Up- and down-regulated genes are shown in red and green, respectively. The Venn diagrams (panel B) represent the total number of genes upregulated (left) and downregulated (right) by at least a factor of 2 under each condition and during growth on glucose in the absence of acetate but at the same growth rate as in the presence of 100 mM acetate (0.35 h^−1^, extrapolated from the data from [40]). Biological functions modulated by the presence of acetate (based on Gene Ontology analysis) are shown in panel C, with the corresponding p-values. The expression levels of central metabolic genes are shown in panel D. The data were obtained from at least three independent biological replicates for each condition.

At the level of the central metabolism (Figure 2D), acetate reduces the expression of all genes that code for the glucose phosphotransferase system (PTS) (*ptsGHI, crr*), without inducing the expression of alternative systems of glucose internalization and phosphorylation (*galP, mglABC*, *glf*, *glk*, *manXYZ*) that could have compensated for reduced PTS activity and the corresponding inhibition of glucose uptake [31, 32]. Expression of upper glycolytic genes remained stable, with the exception of two isoenzymes that were overexpressed in the presence of acetate: *fbaB*, a gluconeogenic enzyme, and *pfkB*, which contributes little to phosphofructokinase activity on glucose (< 10%) [33–36]. In contrast, the expression of most of the lower glycolysis genes (*pgk, gapA, gpmA, eno, pykF, aceE*) was reduced by 15 to 40 % at 100 mM acetate. At this concentration, acetate also inhibits the expression of virtually all TCA cycle genes (*gltA*, *acnAB*, *icd*, *lpd*, *sucABCD*, *sdhABCD*, *fumABC*, *mdh*) by 30 to 67 %. In terms of acetate metabolism, the expression of *pta* and *ackA*, which code for enzymes in the Pta-AckA pathway, remained remarkably stable at all acetate concentrations. Expression of pyruvate oxidase (*poxB*) was increased by a factor of 8 in the presence of acetate, though this promiscuous enzyme contributes to acetate metabolism mainly in the stationary phase and apparently not under our conditions [15, 29, 37, 38]. Expression of acetyl-CoA synthetase (*acs*) – which converts acetate to acetyl-CoA with a high affinity – decreased with increasing acetate concentration, indicating that under glucose excess, the presence of acetate does not activate acetate recycling through the Pta-AckA-Acs cycle [15, 39].

These data confirm our hypothesis that acetate gradually modulates both the flux capacity of *E. coli* to produce acetyl-CoA from glucose and its capacity to utilize acetyl-CoA as a source of energy and of anabolic precursors for growth. Acetate does not appear to influence the flux capacity of the acetate pathway itself, which is also consistent with the proposed model.

### Testing the model

We tested the model by predicting the response of *E. coli* to new perturbations and comparing the results with experimental data (Figure 3).

**Figure 3.**
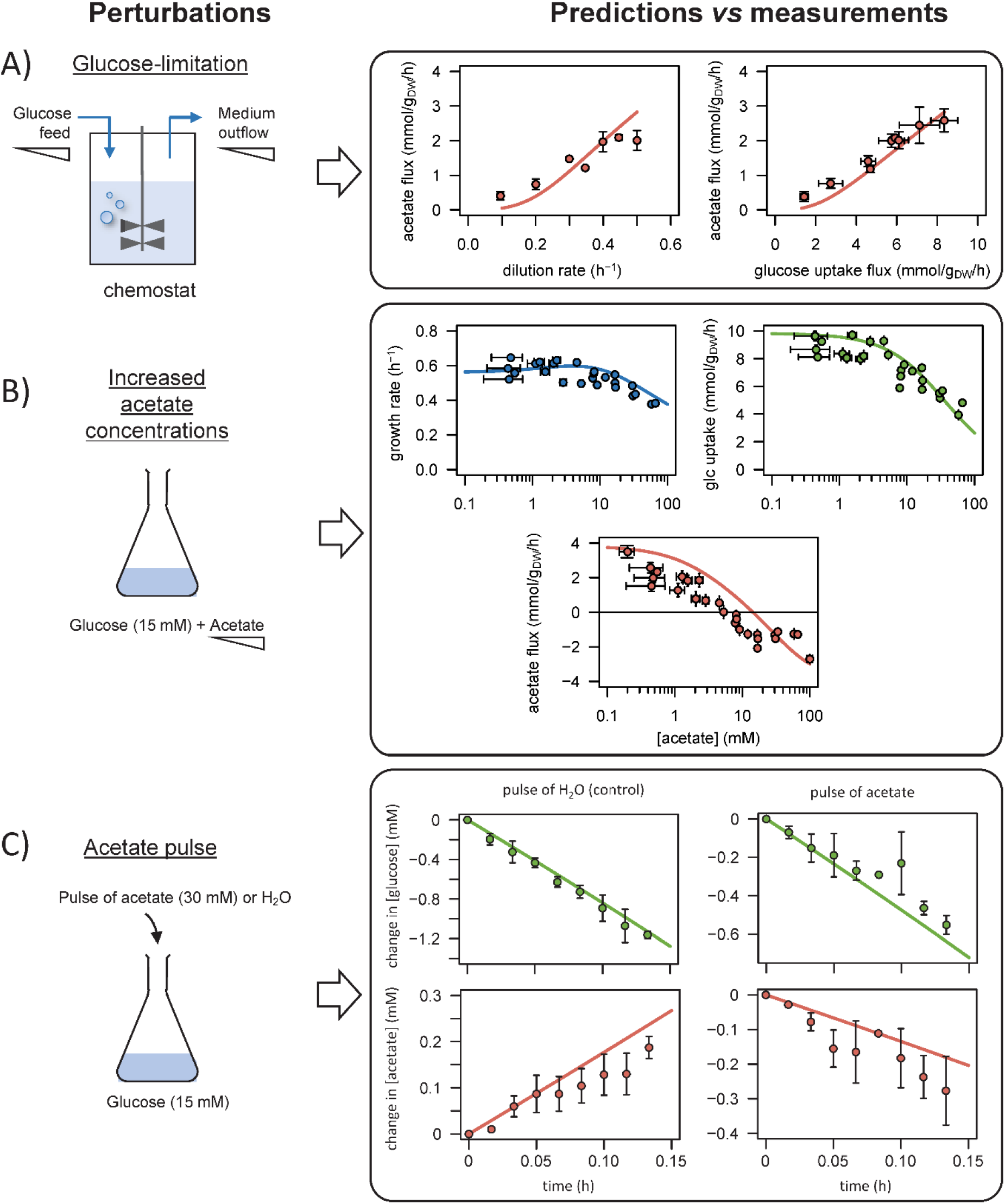
Comparison of model predictions with experimental data. We used the model to simulate i) steady-state glucose and acetate fluxes in glucose-limited chemostat cultures at dilution rates of 0.1–0.5 h^−1^ (panel A), ii) the growth rates and glucose and acetate fluxes during growth on glucose at various acetate concentrations (B), and iii) the time courses of the changes in glucose and acetate concentrations during exponential growth on glucose after a pulse of either acetate or water (C). Model predictions are represented by lines and experimental data are shown as dots (the error bars represent one standard deviation).

First, we checked that the model could reproduce the established growth-rate dependence of acetate production in *E. coli*. We modified the model to simulate glucose-limited chemostat experiments (by adding reactions for glucose feed and medium outflow), and we predicted the steady-state glucose and acetate fluxes at different dilution rates (from 0.1 to 0.5 h^−1^). Since the present model cannot account for the activation of acetyl-CoA synthetase – involved in acetate assimilation – under glucose limitation [39], model simulations were compared to experimental data collected on a Δ*acs* strain [14]. The simulated profiles of acetate production as a function of growth and glucose uptake rates were in excellent agreement with experimental data (Figure 3A), indicating that the model accurately captures the effects of glucose limitation on growth and acetate fluxes.

Second, we evaluated the response of *E. coli* to changes in acetate concentration. We simulated steady-state acetate, glucose and growth fluxes under glucose excess over a broad range of acetate concentrations (between 100 μM and 100 mM) [15]. The growth and glucose uptake rates decreased monotonously with increasing acetate concentrations, in excellent agreement with the experimental data (Figure 3B). The acetate flux profile was also well described by the model, with a progressive decrease of acetate production with increasing acetate concentration, and a reversal of the acetate flux above a threshold concentration of about 10 mM. As observed experimentally, the growth rate was not affected when acetate production was abolished, confirming that acetate production is not required to maintain fast growth nor is it imposed by any intracellular constraint.

Third, we tested the dynamic properties of the model by predicting the response of *E. coli* to sudden variations in the acetate concentration, which is not possible using existing models. In the exponential growth phase on glucose, we simulated the time courses of the changes in the concentrations of glucose and acetate after a pulse of acetate (30 mM) or of water (control experiment), as described in ref. [15]. Here again, the model accurately reproduces the experimental profiles, in particular the rapid reversal of the acetate flux and reduction of glucose uptake after the acetate pulse (Figure 3C).

Overall, the predictions of the model are in agreement with experimental findings. The model accurately predicts the steady-state and dynamic relationships between glucose uptake, growth, and acetate fluxes in *E. coli* over a broad range of glucose and acetate concentrations. Importantly, the model was *not* trained on these data (i.e. they were not used to calibrate the model), so this test is a strong validation of its predictive power.

### Intracellular control of acetate flux is distributed around the acetyl-CoA node

We used a metabolic control analysis of this kinetic model to determine the degree of control exerted by each reaction on acetate flux (Figure 4). Flux control coefficients 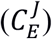 quantify the impact of small changes in the rate of each reaction (typically from a change in the enzyme concentration *E*) on each flux (*J*) [41, 42].

**Figure 4.**
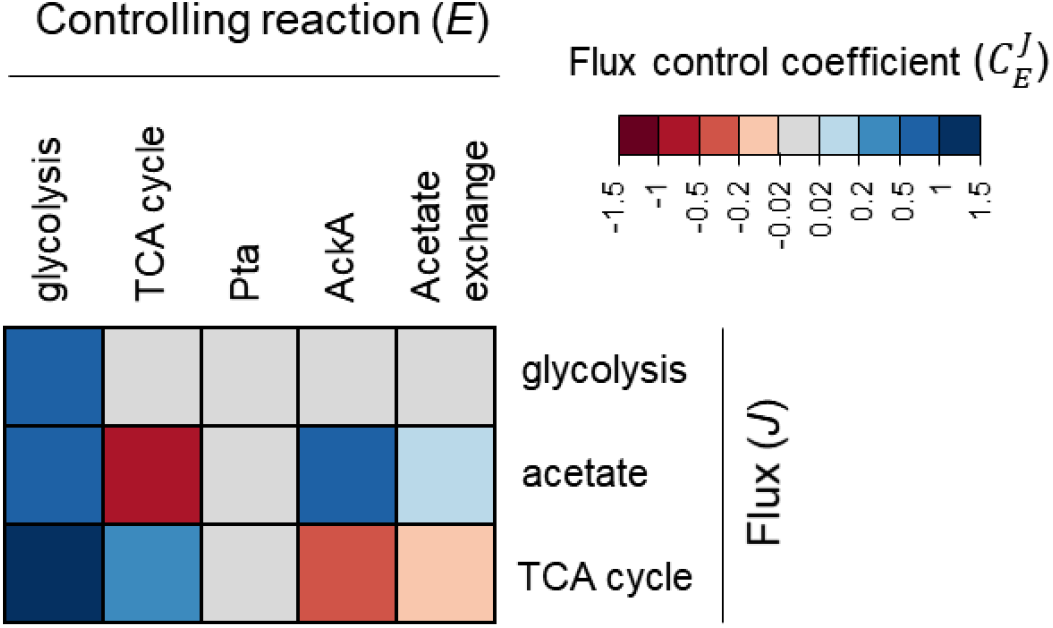
Heatmap of flux control coefficients during growth on glucose (15 mM) and acetate (0.1 mM). Each column represents a controlling reaction (*E*) and each row, a flux (*J*). Red and blue cells represent negative and positive flux control coefficients 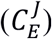, respectively, with darker (lighter) tones indicating stronger (weaker) control.

Metabolic control analysis revealed that rather than being controlled entirely from within the acetate pathway, acetate flux is controlled to a similar degree by the acetate pathway (with a flux control coefficient of 0.76), glycolysis (0.88), and the TCA cycle (−0.64). The balance between acetyl-CoA production and demand (i.e. between glycolytic and TCA fluxes) therefore has a strong effect on acetate production. As expected, this control is positive for glycolysis (since it produces acetyl-CoA) and negative for the TCA cycle (since it consumes acetyl-CoA). Still, acetate flux is controlled to a large extent from within the acetate pathway, mainly by AckA (0.69), with a small contribution from acetate transport (0.07). Overall, acetate flux control is distributed around the acetyl-CoA node.

### Intracellular flux control patterns shift with the acetate concentration

Since control of acetate flux is distributed around the acetyl-CoA node, we tested how flux changes around this node might affect its control properties. Given that the concentration of acetate is a major determinant of acetate flux (Figure 3), we performed metabolic control analyses for a broad range of acetate concentrations (from 0.1 to 100 mM).

The control exerted by the acetate pathway on acetate flux decreases non-linearly as the acetate concentration increases (Figure 5A), highlighting a progressive shift of the control from inside to outside the acetate pathway. This is contrary to the classical behaviour in which the control exerted by a metabolic pathway increases as it becomes saturated, i.e. where the flux reaches the capacity of the uptake pathway at high substrate concentrations [43, 44]. The control exerted by the acetate pathway gradually shifts towards the glycolytic and TCA pathways (Figure 5B, C). Discontinuous control patterns are observed for the latter pathways at a concentration of about 10 mM, i.e. around the concentration threshold at which the acetate flux switches from production to consumption and is zero [15]. The sum of the control coefficients of the glycolytic and TCA blocks, which represent the overall control exerted by *E. coli* metabolism on the acetate flux (Figure 5D), increases with the acetate concentration and compensates for the decrease in control from the acetate pathway. The ratio of the control coefficients of glycolysis and the TCA cycle show that control shifts from the glycolytic pathway to the TCA cycle as the acetate concentration increases (Figure 5E).

**Figure 5.**
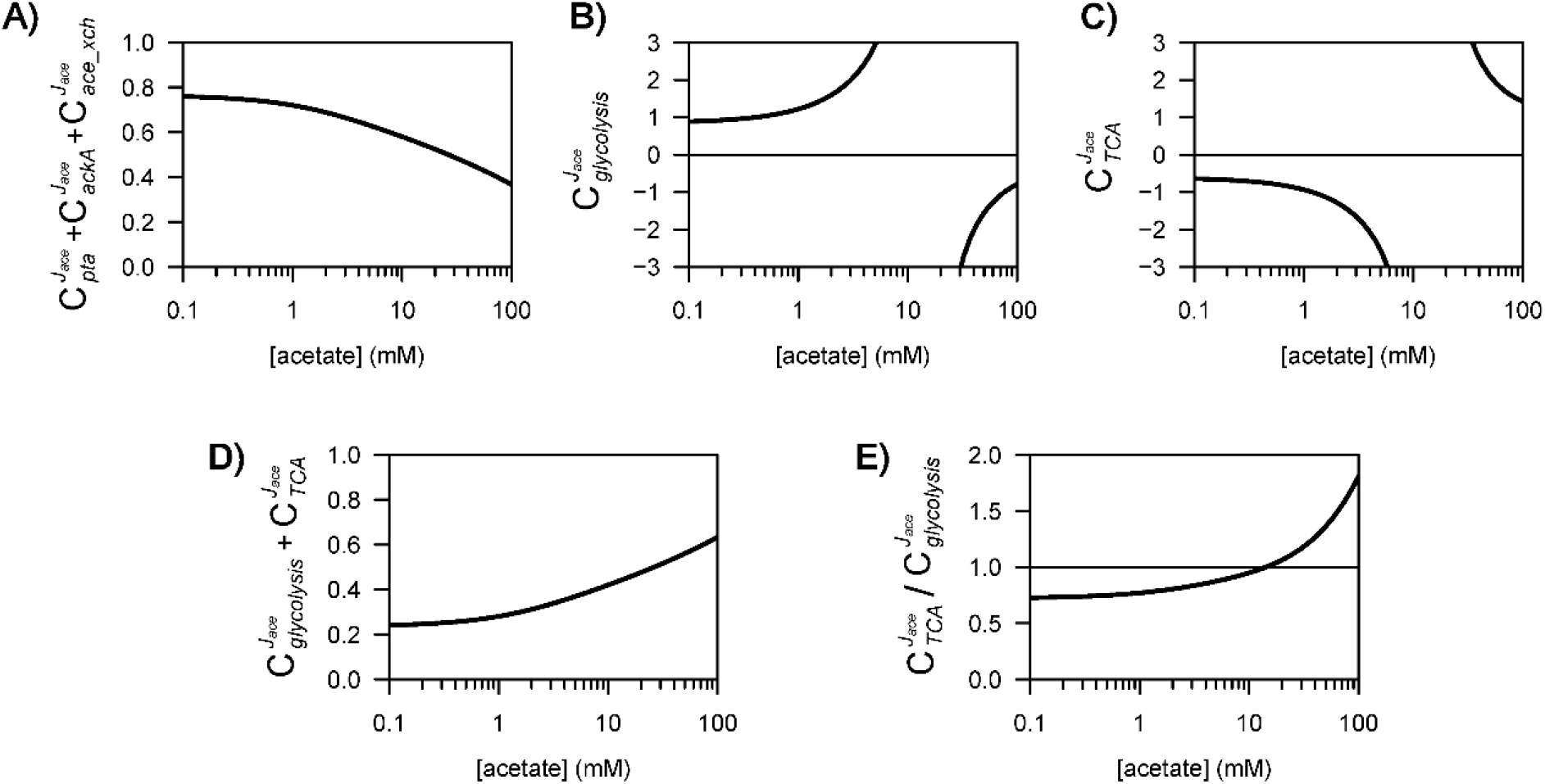
Control of acetate flux over a broad range of acetate concentrations.

Overall, these results reveal how intracellular flux control patterns are strongly modulated by the extracellular concentration of acetate, with the control exerted by the acetate pathway being transferred progressively to the glycolytic and TCA pathways as the acetate concentration increases.

### Regulation of E. coli metabolism by acetate

Metabolic control analysis identified the controlling steps, i.e. the reactions that alter the acetate flux if their rates are modified (e.g. through changes in enzyme concentrations). However, more information is needed to understand the mechanisms involved in flux responses to perturbations of an external parameter such as the acetate concentration. Indeed, while a metabolic reaction may exert some control on a given flux, this does not necessarily mean that it is involved in the observed flux response. For instance, AckA exerts some *control* on the acetate flux, but its expression appears to be constant at all acetate concentrations (Figure 2), hence it does not *regulate* the acetate flux at the transcriptional level.

To identify the regulatory routes that are actually involved in the response of *E. coli* to changes in acetate concentration, we used the concept of response coefficients 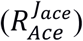, which express the dependence of a system variable (here the acetate flux, *J_ace_*) on an external effector (the concentration of acetate). The partitioned response relationship [41, 45] allows the flux response to a perturbation in acetate concentration channeled through a given reaction *i* 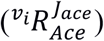 to be quantified, as detailed in the *Materials and methods* section. Since acetate regulates more than one reaction, the partitioned response coefficients provide a quantitative understanding of the different routes through which the acetate flux is regulated by the acetate concentration. We calculated the partitioned response coefficients of the acetate flux to the acetate concentration via i) the acetate pathway, which represents the contribution of direct metabolic regulation, and via ii) the glycolytic and iii) the TCA pathways, where acetate acts indirectly by modulating the flux capacity (Figure 6A). As expected, regulation is minimal when acetate does not significantly modulate the acetate flux, i.e. at low and high acetate concentrations, and is maximal at concentrations that strongly modulate its flux (Figure 6B-D). Setting the response threshold of 0.1 – i.e. that a relative change of *x* % in the acetate concentration should lead to a relative flux response of at least (0.1 × *x*) % via the regulatory route considered –, we determined that acetate acts as a regulator over a range of concentrations spanning three orders of magnitude (between 0.2 and 100 mM).

**Figure 6.**
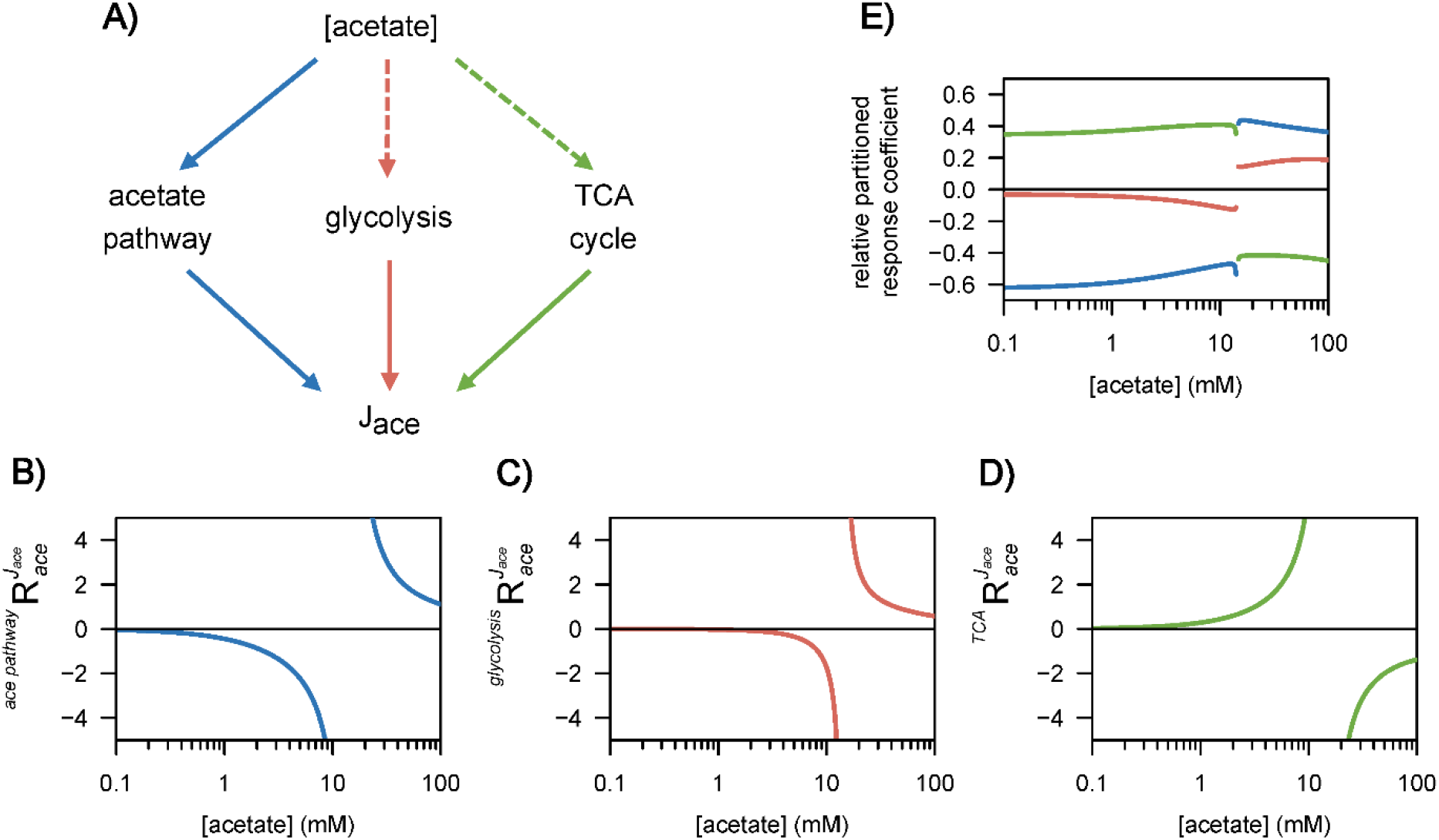
Regulation of acetate flux in *E. coli*. The different routes through which acetate flux can be regulated by the acetate concentration are shown in panel A. Dotted lines represent indirect (hierarchical) regulation, and straight lines represent direct (metabolic) regulation. The strengths of the three regulatory routes are respectively shown in panels B–D, and their relative contributions are shown in panel E.

Based on the regulatory strength of each interaction, we determined the relative contribution of each interaction to the observed flux response (Figure 6E), revealing two distinct modes of regulation that depend on the acetate concentration and remain remarkably stable over a broad range of concentrations. At low acetate concentrations (< 10 mM), the main regulatory route (50–60%) is local metabolic regulation via the acetate pathway, with the remaining flux response being accounted for by hierarchical regulation through the TCA cycle (35–40%) and glycolysis (5-10%). This regulatory pattern remains stable before changing abruptly at ~10 mM when the acetate flux reverses. Above this threshold, direct regulation by acetate accounts for about 40% of the observed flux response, glycolysis, about 20%, and the TCA cycle, roughly 40%. This pattern then remains stable up to 100 mM.

These results provide a comprehensive and quantitative understanding of acetate flux regulation by acetate through the coordinated effects of direct and indirect mechanisms, with distinct regulatory programs at low and high acetate concentrations.

## Discussion

Two independent theories have been proposed to explain acetate overflow in *E. coli*, but neither global regulation of central metabolic pathways nor local control of the acetate pathway can account for all observations. Using a systems biology strategy, we reconcile these two theories, clarify how acetate flux is controlled and regulated, and establish the pivotal role acetate plays as a global regulator of glucose metabolism in *E. coli*.

The kinetic model of *E. coli* metabolism developed in this work links acetate metabolism with glucose uptake and growth. This model successfully predicts the steady-state and dynamic responses of *E. coli* to a broad range of perturbations. The model also suggested the existence of a global regulatory program whereby the extracellular concentration of acetate determines the flux capacity of the glycolytic and TCA pathways, whose inhibition was required to explain the dynamic profiles obtained in ^13^C-labeling experiments. Transcriptomics experiments confirmed that acetate controls the flux capacity of these two pathways at the transcriptional level by modulating the expression of most of their genes. This is the most efficient way for cells to adjust their fluxes while maintaining metabolite homeostasis [41, 42]. The metabolic response of *E. coli* to changes in acetate concentration involves a global reorganization of its metabolism from the transcriptional to the flux levels, contrary to what has been suggested [29] and providing a mechanistic rationale for the reported “toxicity” of acetate. The coordinated regulation of the glycolytic and TCA pathways cannot be explained by the actions of known transcriptional regulators, suggesting that these regulators act in concert or that there are additional regulators present. It is likely that acetate also regulates *E. coli* metabolism at the post-translational level (e.g. via the carbon storage regulator system [46] or acetyl-phosphate [47, 48]) and at the metabolic level (e.g. via the control of glucose uptake by α-ketoglutarate [49] or of glycolysis by ATP [50]).

Our results confirm that the direction of the acetate flux is controlled thermodynamically and is fundamentally determined by the extracellular concentration of acetate. Moreover, we found that intracellular control of the acetate flux is distributed around the acetyl-CoA node, i.e. between the glycolytic, TCA, and acetate pathways, such that none of the three processes can be regarded as the sole kinetic bottleneck. The intracellular flux control pattern is determined by the concentration of acetate itself, with a progressive shift of control from the acetate pathway to the rest of the metabolism when the acetate concentration increases. This means that metabolic engineering interventions aiming at minimizing acetate production in biotechnology should rebalance the flux capacity of both the glycolytic and the TCA pathways, possibly via global metabolic regulators, while accounting for the concentration-dependent effect of acetate on intracellular control patterns. The proposed model may thus be used to guide rational design of optimized strains for synthetic biology applications.

This study also identifies the regulatory scheme involved in *E. coli’s* response to changes in acetate concentration, which largely determines the acetate flux itself as well as the cell’s physiology. We demonstrate that this scheme involves direct metabolic regulation of the acetate pathway in combination with transcriptional regulation of the glycolytic and TCA pathways, and we show how this equal contribution of direct and indirect regulatory mechanisms explains the diverse patterns of acetate flux observed and forms part of *E. coli’s* global physiological response to changing acetate concentrations. The distinct regulatory patterns at high and low acetate concentrations also determine the metabolic status of acetate as a co-substrate or by-product of glucose metabolism. This work shows that the regulation of acetate flux is far more subtle than previously considered.

These results call for a reconsideration of the role of acetate, and more generally of overflow metabolism, in *E. coli*. Acetate should not be considered a terminal, wasteful product of glucose metabolism since it may equally be a co-substrate of glucose. Moreover, it is also a key regulator that induces a global remodeling of *E. coli* physiology and modulates several cellular functions (motility, biofilm formation, transport, metabolism, translation). Acetate regulates *E. coli* metabolism over a range of concentrations spanning three orders of magnitude, and its regulatory action involves a combination of direct and indirect mechanisms. In the laboratory, *E. coli* is typically grown on glucose at low acetate concentrations, which enhances its production. In the intestine however, *E. coli’s* environmental niche, the concentration of acetate is high (between 30 and 100 mM) [23, 24] and well above the ~10 mM threshold at which the acetate flux reverses. These levels of acetate should thus favour its co-utilization with available sugars, suggesting that acetogenic species may be an important source of nutrients for *E. coli*. Besides its importance in terms of understanding cross-feeding relationships, this work also stresses the need to further investigate the regulatory role of acetate (and of acetogenic species) in the intestine. This will require a quantitative characterization of the available nutrients and of their dynamics in this particularly complex environment.

Other (micro)organisms than *E. coli* are similarly capable of co-consuming glucose with its metabolic by-products, even when glucose is abundant. *Saccharomyces cerevisiae* can co-consume ethanol and glucose [51], and mammalian cells co-assimilate lactate with glucose [52–54]. These similarities, which have important implications for cell energy and redox balance, indicate that overflow metabolism should be considered a reversible process and point to the existence of a universal phenomenon similar to the one described here for acetate in *E. coli*. The quantitative systems biology approach developed in this work can be used as a guide for future investigations of overflow fluxes, their regulation, and their biological implications in *E. coli* and others (micro)organisms.

## Materials and Methods

### Strain and conditions

*Escherichia coli* K-12 MG1655 was grown in M9 medium [55] complemented with 15 mM glucose. Sodium acetate (prepared in a 1.6 M solution at pH 7.0) was added up to the required concentration. The cells were grown in shake flasks at 37 °C and 200 rpm, in 50 mL of medium. For the isotope labelling experiments, unlabelled glucose was replaced by U-^13^C_6_-glucose (Eurisotop, France). Growth was monitored by measuring the optical density (OD) at 600 nm using a Genesys 6 spectrophotometer (Thermo, USA), and a conversion factor of 0.37 g_DW_/L/OD unit [46] was used to determine the biomass concentration.

### Transcriptomics experiments

Cells were grown in flasks in M9 minimal media with 15 mM glucose and 0, 10, 50 or 100 mM acetate. In mid-exponential growth phase (OD_600nm_ = 1), 4 mL of each culture was centrifuged for 90 s at 14000 rpm, the supernatant was discarded, and the pellets were immediately frozen in liquid nitrogen. Total RNA was extracted using a Qiagen RNAeasy MiniKit and quantified using a Nanodrop spectrophotometer. Double-stranded complementary DNA (cDNA) synthesis and array processing were performed by One-Color Microarray-Based gene Expression Analysis (Agilent Technologies). The images were analysed with the software DEVA (v1.2.1). All array procedures were performed using the GeT-Biopuces platform (http://get.genotoul.fr). For each data set, the log2 intensities obtained in the presence of acetate were divided by the log2 intensities obtained in the absence of acetate. These ratios were then normalized by the log2 median intensity. Genes whose expression level differed by a factor of 2 or more between the two conditions were used for further analysis. Gene ontology analyses were performed using Ecocyc (https://ecocyc.org). The transcriptomics data can be downloaded from the ArrayExpress database (www.ebi.ac.uk/arrayexpress) under accession number E-MTAB-9086. Theoretical expression data at a given growth rate were obtained by extrapolating the data from ref. [40].

### Metabolomics experiments

Extracellular concentrations of labelled and unlabelled glucose and acetate were quantified in 150 μL of filtered broth (0.2 μm, Sartorius, Germany) by 1D ^1^H-NMR on a Bruker Ascend 800 MHz spectrometer equipped with a 5 mm QCI cryoprobe (Bruker, Germany), as detailed previously [56].

### Model construction and analysis

The models are briefly described in this section, and additional information can be found in the Supplementary Information (Supplementary File 1). All models are provided in SBML and COPASI formats in the Supplementary Information (Supplementary File 2) and at https://github.com/MetaSys-LISBP/acetate_regulation. The kinetic model has also been deposited in the Biomodels database (https://www.ebi.ac.uk/biomodels) [57] with the identifier MODEL2005050001. Model analysis was performed using COPASI [58] (v4.27) with the *CoRC* package (COPASI R Connector v0.6.1, https://github.com/jpahle/CoRC) in R (v3.6.1, https://www.r-project.org). The scripts used to perform the simulations, to analyse the models and to generate the figures are provided in the Supplementary Information (Supplementary file 2) and at https://github.com/MetaSys-LISBP/acetate_regulation to ensure reproducibility and reusability.

#### Model construction

The model contains 6 reactions, 6 species, and 2 compartments (the environment and the cell). Glycolysis (which produces acetyl-CoA from glucose) and the TCA cycle (which utilizes acetyl-CoA to produce biomass) were modelled using irreversible Michaelis-Menten kinetics. Growth rates were calculated from the flux of the TCA cycle assuming a constant biomass production yield from acetyl-CoA, in keeping with observations [15, 29]. Acetate exchange between the cell and its environment was modelled as a saturable process using reversible Michaelis-Menten kinetics [26], and the Pta-AckA pathway was modelled using the detailed kinetics of the Pta and AckA enzymes used in previous models [15, 26, 27]. As detailed in the *Results* section, we constructed four different versions of this model, i.e. with or without (non-competitive) inhibition of the glycolytic and/or TCA cycle pathways by acetate. Finally, these models were extended with isotopic equations for parameter estimation, as detailed in refs. [15, 28].

#### Parameter estimation

The values of 19 of the 29 parameters were taken directly from the literature (Supplementary File 1). Parameters whose values are not available from elsewhere, which do not have a real biochemical value, or for which biochemical measurements are generally not representative of intracellular conditions (e.g. Vmax) were estimated to optimally reproduce 152 experimental data points obtained from *E. coli* K-12 MG1655 grown on ^13^C-glucose (15 mM) plus ^12^C-acetate (1, 10 or 30 mM). These data included time-course concentrations of biomass, glucose and acetate and ^13^C-labeling of acetate. The parameters were estimated by minimizing the objective function *f* defined as the weighted sum of squared errors:

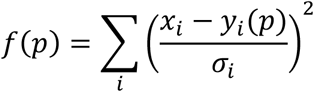

where *x_i_* is the experimental value of data point *i*, with and experimental standard deviation *σ_i_*, and *y_i_*(*p*) is the corresponding simulated value. The objective function *f* was minimized with the particle swarm optimization algorithm (2000 iterations with a swarm size of 50). The experimental and fitted data are shown in Figure 1 and provided in the Supplementary Information (Supplementary File 2). The values of all the parameters (and the corresponding references for the values taken from the literature) are also listed in the Supplementary Information (Supplementary File 1).

#### Goodness-of-fit analysis

We used *χ*^2^ statistical tests to assess the goodness-of-fit of each model and determine whether they described the data with sufficient accuracy. The minimized variance-weighted sum of squared residuals (SSR) is a stochastic variable with a *χ*^2^ distribution. The acceptable threshold for SSR values is *χ*^2^(*α*, *d*), where *d* represents the number of degrees of freedom and is equal to the number of fitted measurements *n* minus the number of estimated independent parameters *p*. The parameter α was set to 0.95 to define a 95 % confidence threshold and models with SSRs above this threshold were rejected since they cannot accurately reproduce the experimental data.

#### Model validation

A total of 170 experimental data (extracellular fluxes and concentrations) were obtained from the literature for *E. coli* K-12 MG1655 and its close derivative strain BW25113 grown on glucose with or without acetate [14, 15, 29], as detailed in the *Results* section. The good agreement between simulated and experimental validation data (Figure 3) indicates that the model yielded accurate predictions of the steady-state and dynamic responses of *E. coli* to glucose and acetate perturbations.

#### Metabolic control and regulation analyses

Scaled flux control coefficients 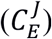, which represent the fractional change in the steady-state flux *J* in response to a fractional change in the rate of the reaction *E* [41, 42, 45], were calculated as follows:

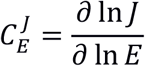

Similarly, we defined the response coefficient 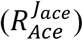 which represents the dependence of the acetate flux (*J_ace_*) on the extracellular concentration of acetate [41, 45]:

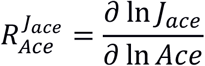

The partitioned response relationship [41, 45] was used to quantify the flux response 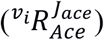 to a change in acetate concentration channeled through reaction *i*. The effect of the acetate concentration on the rate of reaction *i* (*v_i_*) is described by the elasticity coefficient 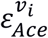, and the resulting change in *v_i_* then propagates through the system depending on the control exerted by reaction *i* on the acetate flux 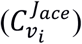:

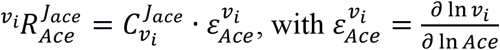

## Acknowledgements

The authors thank MetaboHub-MetaToul (Metabolomics & Fluxomics facilities, Toulouse, France, http://www.metatoul.fr), which is part of the French National Infrastructure for Metabolomics and Fluxomics (www.metabohub.fr), funded by the ANR (MetaboHUB-ANR-11-INBS-0010), for access to NMR and MS facilities. JCP is grateful for funding from INSEMR for his temporary full-time researcher position. The authors are grateful for Vincent Pierunek’s help with the isotope labelling experiments, and wish to thank the following INSA Toulouse students for help with the transcriptomics experiments: Leidy Caraballo, Xavier Caron, Pauline Chanut, Sarah Guiziou, Ngoc Thu Hang Pham, Diane Barbay, Céline Ben Hassen, Thomas Cerutti, Cécile Roland, Sarah Srour, Audrey Baylet, Mathilde Beraud, Claudie Bosc, Lilas Courtot, Violaine Dolfo, Anna Kaci, Manon Chevallot-Beroux, Sarah Colom, Fanny Leclerc, Zihan Liao, Grégoire Quinet and Mélina Vaurs.

## Contributions

Funding acquisition: PM, JCP, FL; Project administration: PM; Conceptualization: PM; Investigation: PM, BE, SUJ; Methodology: PM, BE; Software: PM; Formal Analysis: PM, BE; Visualization: PM, BE; Writing – original draft: PM; Writing – review & editing: PM, BE, SUJ, JCP, FL; Supervision: PM.

