## Supplementary file 1 for "Control and regulation of acetate overflow in *Escherichia coli*"

### Kinetic model of glucose and acetate metabolisms of *Escherichia coli*

-

Documentation

*Corresponding author

#### 1. Model overview

The models developed in this work represent glucose and acetate metabolisms of the bacterium *Escherichia coli*. It should be noted that, when developing a model, certain criteria must be decided, such as the level of detail and the boundaries within which the model can be expected to be valid. The current model simulates the metabolic functioning of *E. coli* K-12 MG1655 during growth on glucose (plus acetate) under aerobic condition. It may allow simulation of other scenarios by changing the parameter values to reflect the altered conditions, and/or implementing additional pathways known to be active in the other scenarios.

All models are available in SBML and COPASI formats in Supplementary data and at [https://github.com/MetaSys-LISBP/acetate_regulation](https://github.com/MetaSys-LISBP/acetate_regulation). The kinetic model can also be downloaded from the Biomodels database ([http://www.ebi.ac.uk/biomodels](http://www.ebi.ac.uk/biomodels)) with identifier MODEL2005050001.

This model comprises 2 compartments (the environment and the cell), 6 species and 6 reactions that represent the following processes (Figure 1):

- glucose uptake and glycolysis (reaction *glycolysis*)
- tricarboxylic acids cycle (reaction *TCA_cycle*)
- acetate metabolism (reactions *Pta*, *AckA* and *acetate*_exchange)
- biomass synthesis (coupled to reaction *TCA_cycle*)

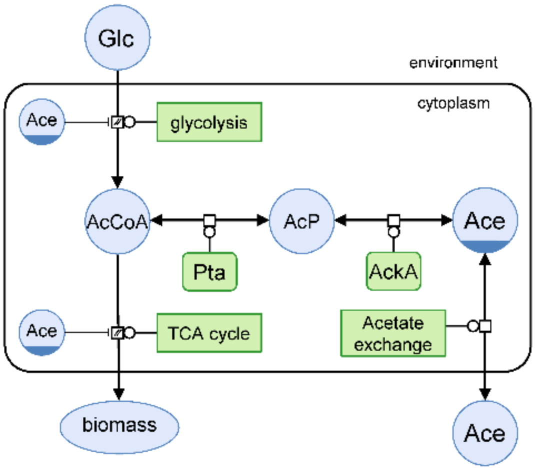

**Figure 1.** Metabolic network of *E. coli* glucose and acetate metabolisms. Metabolites are shown in blue, and enzymes/pathways are shown in green. The diagram follows the conventions of the Systems Biology Graphical Notation process description [1].

#### 2. Model units

Model units are millimole (mmol) for amounts, litre (L) for volumes, and hour (h) for time. Amount of biomass is expressed as gram dry weight ($g_{DW}$).

#### 3. Cell volume

Cell volume (V) was set to 1.77 × $10^{-3}$ L/$g_{DW}$ [2].

#### 4. Reactions

The reactions included in the model are listed in the table below. When the rate law was taken from the literature, the corresponding reference is given.

| ***Name*** | ***Reaction*** | ***Rate law***[a] | ***Comment*** |
|---|---|---|---|
| Glucose_feed | ∅ → GLC | Constant flux | Glucose inflow and medium outflow to simulate chemostat experiments |
| Acetate_outflow | $ACE_{env}$ → ∅ | MA | |
| Biomass_outflow | X → ∅ | MA | |
| Glucose_outflow | GLC → ∅ | MA | |
| Glycolysis | GLC → 1.4 * ACCOA | IMM | Stoichiometric coefficient taken from [3] |
| TCA_cycle | ACCOA → ∅ | IMM | - |
| Pta | ACCOA ↔ ACP | RMM | Rate law from [4-6] |
| AckA | ACP ↔ $ACE_{cell}$ | RMM | Rate law from [4-6] |
| Acetate_exchange | $ACE_{cell}$ ↔ $ACE_{env}$ | RMM | Rate law from [5] |
| Growth | X → 2 * X | MA | Rate calculated from the TCA cycle flux, assuming a constant biomass yield [6, 7] |

[a]MA: Mass action; RMM: Reversible Michaelis-Menten; IMM: Irreversible Michaelis-Menten.

#### 5. ODEs system

The differential equations, which describe the progression of the variables over time as a function of the system's rates, balance the concentrations of extracellular (biomass, glucose and acetate) and intracellular (acetate, acetyl-CoA and acetyl-phosphate) species.

<u>Extracellular species:</u>

$$\frac{dGLC}{dt} = v_{glycolysis} \cdot X \cdot \frac{V_{cell}}{V_{env}} \; [+v_{feed} - D \cdot GLC]$$

$$\frac{dACE_{env}}{dt} = v_{acetate\_exchange} \cdot X \cdot \frac{V_{cell}}{V_{env}} \; [-D \cdot ACE_{env}]$$

$$\frac{dX}{dt} = X \cdot v_{growth} \; [-D \cdot X]$$

Note: terms within square brackets are required only to simulate chemostat experiments (at dilution rate $D$ and with a glucose feed defined by $v_{feed}$).

Intracellular species:

$$\frac{dACCOA}{dt} = 1.4 \cdot v_{glycolysis} - v_{TCA_{cycle}} - v_{Pta}$$

$$\frac{dACP}{dt} = v_{Pta} - v_{AckA}$$

$$\frac{dACE_{cell}}{dt} = v_{AckA} - v_{acetate\_exchange}$$

#### 6. Rate laws

This section contains the rate laws for each reaction. Terms within square brackets refers to inhibition of glycolysis and TCA cycle by acetate, as detailed in the publication.

$$v_{glycolysis} = \frac{Vmax_{glycolysis} \cdot GLC}{GLC + Km\_GLC} \quad \left[ \cdot \frac{1}{1 + \frac{ACE_{env}}{Ki\_ACE}} \right]$$

$$v_{TCA\_cycle} = \frac{Vmax_{TCA\_cycle} \cdot ACCOA}{ACCOA + Km\_ACCOA} \quad \left[ \cdot \frac{1}{1 + \frac{ACE_{env}}{Ki\_ACE}} \right]$$

$$v_{AckA} = \frac{\frac{Vmax_{AckA} \cdot \left( ACP \cdot ADP - \frac{ACE_{cell} \cdot ATP}{Keq} \right)}{Km\_ACP \cdot Km\_ADP}}{\left( 1 + \frac{ACP}{Km\_ACP} + \frac{ACE_{cell}}{Km\_ACE} \right) \cdot \left( 1 + \frac{ADP}{Km\_ADP} + \frac{ATP}{Km\_ATP} \right)}$$

$$v_{Pta} = \frac{\frac{Vmax_{Pta} \cdot \left( ACCOA \cdot P - \frac{ACP \cdot COA}{Keq} \right)}{Km\_ACCOA \cdot Km\_P}}{1 + \frac{ACCOA}{Km\_ACCOA} + \frac{P}{Ki\_P} + \frac{ACP}{Ki\_ACP} + \frac{COA}{Km\_COA} + \frac{ACCOA \cdot P}{Km\_ACCOA \cdot Km\_P} + \frac{ACP \cdot COA}{Km\_ACP \cdot Km\_COA}}$$

$$v_{Acetate\_exchange} = \frac{\frac{Vmax_{AckA} \cdot \left( Ace_{cell} - \frac{ACE_{env}}{Keq} \right)}{Km\_ACE \cdot Km\_ACE}}{1 + \frac{Ace_{cell}}{Km\_ACE} + \frac{ACE_{env}}{Km\_ACE}}$$

$$v_{Growth} = v_{TCA\_cycle} \cdot Y$$

#### 7. Concentrations of cofactors

Concentrations of cofactors were taken from a published kinetic model of the Pta-AckA pathway [6] and are listed in the table below.

| Specie | Intracellular concentration (mM) |
|---|---|
| COA | 1.22 |
| ADP | 0.61 |
| ATP | 2.40 |
| Phosphate | 10 |

We evaluated if increasing the extracellular concentration of acetate affects the intracellular concentration of these (unbalanced) cofactors. *E. coli* was grown on glucose (15 mM) plus acetate (0, 10, 50 or 100 mM), and intracellular concentrations of ADP, ATP and CoA were quantified by LC-MS using the differential method and the IDMS approach [8]. In mid-exponential growth phase (OD ≈ 1), 120 µL (for ADP and ATP) or 1 mL (for CoA) of broth and filtered medium were rapidly sprayed into precooled centrifuged tubes maintained at –20 °C and containing 5 mL of quenching and extraction solutions (80 % methanol / 20 % formic acid 125 mM in water for CoA, and 40 % acetonitrile / 40 % methanol / 20 % formic acid 50 mM in water for ADP and ATP), and 100 µL of a fully $^{13}$C-labeled cellular extract was added as internal standard. Samples were kept at -20 °C during 1 h, homogenized using a vortex, and centrifuged (12 000 g at -20 °C for 5 min using a Sigma 3-18K, Sigma). Supernatants were evaporated (SC110A SpeedVac Plus, ThermoFisher) and resuspended in 120 µL of resuspension solution (milliQ water for ADP and ATP, 98 % 25 mM ammonium formate in water (at pH 3.0) / 2% methanol for CoA). Cell extracts were stored at -80 °C until analysis. ADP and ATP were quantified as detailed in [3]. CoA was quantified using a Thermo Scientific Vanquish Focused UHPLC Plus system, coupled to a Thermo Scientific Q Exactive Plus mass spectrometer (Thermo Fisher Scientific). Separation was performed on a Phenomenex Kinetex column (100 mm × 3.0 mm; 1.7 µm) at 25 °C and with a flow rate of 0.35 mL/min. Solvent A was 25 mM ammonium formate in water (pH 8.0) and solvent B was methanol. The following gradient of B was applied: 0-2 min, 2%; 2-23 min, 25-95%; 23-28 min, 95%. Injection volume was 5 µL. MS analysis was performed by Full Scan mode at a resolution of 70 000 (from *m/z* 300 to 1500), using the positive electrospray ionization (ESI) mode. Spray voltage was -4.5 kV, capillary temperature was 320 °C, desolvation temperature was 350 °C, and maximum injection time was 50 ms. Nitrogen was used as sheath gas (pressure 55 units) and auxiliary gas (pressure 10 units). The automatic gain control (AGC) was set at $10^6$. Data acquisition and processing was performed using Thermo Scientific Xcalibur software.

Quantification results (Figure 2), which are in good agreement with literature values, indicate that these pools remained stable under a broad range of acetate concentrations (between 0 and 100 mM). This was also the case for the ATP/ADP ratio.

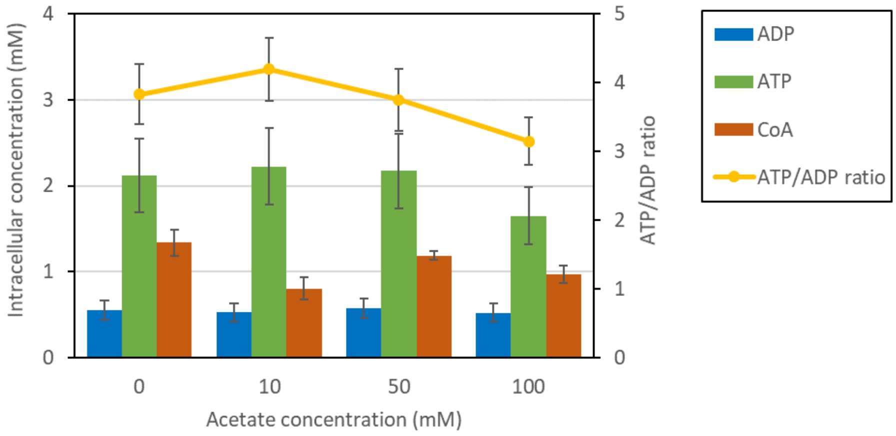

**Figure 2.** Intracellular concentration of ADP, ATP and CoA, and ATP/ADP ratio, during growth on glucose (15 mM) plus acetate (0, 10, 50 or 100 mM).

#### 8. Extension with isotopic equations

The kinetic model was extended with isotopic equations as detailed in [6, 9]. Briefly, all reactions (except biomass synthesis) were considered separately for unlabeled and labeled metabolites. Rate laws of reversible reactions were decomposed into their forward and reverse components to account for the bidirectional isotope exchange that arise from reversibility and significantly influences the distribution of isotopes through the network.

For instance, the isotopically extended balance of acetyl-CoA corresponds to:

$$\frac{dACCOA_0}{dt} = \frac{GLC_0}{GLC_0 + GLC_1} \cdot 1.4 \cdot v_{glycolysis} + \frac{ACP_0}{ACP_0 + ACP_1} \cdot v_{Pta}^{reverse} - \frac{ACCOA_0}{ACCOA_0 + ACCOA_1} \cdot \left(v_{TCA_{cycle}} + v_{Pta}^{forward}\right)$$

$$\frac{dACCOA_1}{dt} = \frac{GLC_1}{GLC_0 + GLC_1} \cdot 1.4 \cdot v_{glycolysis} + \frac{ACP_1}{ACP_0 + ACP_1} \cdot v_{Pta}^{reverse} - \frac{ACCOA_1}{ACCOA_0 + ACCOA_1} \cdot \left(v_{TCA_{cycle}} + v_{Pta}^{forward}\right)$$

where subscripts 0 and 1 refers to the unlabeled and labeled metabolite, respectively.

#### 9. Model calibration

Unknown parameters were estimated by fitting experimental data, as detailed in the publication. Values obtained from the best fit are provided below.

| *Reaction* | *Parameter* | *Value* | *Source* |
|---|---|---|---|
| AckA | Keq | 174 | [4-6] |
| | KmACE | 7 | [5, 6, 10] |
| | KmACP | 0.16 | [5, 6, 10] |
| | KmADP | 0.5 | [5, 6, 10] |
| | KmATP | 0.07 | [5, 6, 10] |
| | Vmax | 336000 | Estimated |
| Pta | Keq | 0.005 | [5, 6] |
| | KmACCOA | 0.2 | [5, 6, 11] |
| | KiACP | 0.2 | [5, 6, 11] |
| | KmCOA | 0.029 | [5, 6, 11] |
| | KiP | 13.5 | [5, 6, 11] |
| | KmACP | 0.7 | [5, 6, 11] |
| | KmP | 6.1 | [5, 6] |
| | Vmax | 976500 | Estimated |
| Glycolysis | KmGLC | 0.020 | [12] |
| | Vmax | 5557 | Estimated |
| | Ki | 36.7 | Estimated |
| TCA_cycle | KmACCOA | 24.8 | Estimated |
| | Vmax | 741000 | Estimated |
| | Ki | 2.3 | Estimated |
| Growth | Y | 0.0000998 | Estimated |
| Acetate_exchange | Keq | 1 | Intracellular and extracellular concentrations equilibrate over time |
| | Vmax | 480035 | Estimated |
| | Km | 33.2 | Estimated |
