## Supplementary file 2 for "Control and regulation of acetate overflow in *Escherichia coli*": README.md

# Kinetic modeling of glucose and acetate metabolisms in E. coli - Millard et al., 2020.

## Overview

This R script performs all in silico analyses detailed in the following publication:

> Control and regulation of acetate overflow in *Escherichia coli*
> 
> Millard et al., 2020, bioRxiv preprint

All models and experimental data are available in COPASI format in the directory `/model/cps/`. The final kinetic model is also available in SBML format in the 
directory `/model/sbml/` and from the Biomodels database (http://www.ebi.ac.uk/biomodels/) under identifier MODEL2005050001.

Details on the calculations can be found in the publication and in the script `run_analysis.R`.

## Dependencies

Some R packages are required.

`RColorBrewer` and `gplots` can be installed
by running the following command in an R console:

```bash
install.packages(c("RColorBrewer", "gplots"))
```

`CoRC` can be installed
using the following command:

```bash
install.packages("remotes")
library(remotes)
remotes::install_github("jpahle/CoRC")
library(CoRC)
CoRC::getCopasi()
```

Additional information on CoRC installation and usage are available from the CoRC repository (https://github.com/jpahle/CoRC).

## Usage

To run all analyses detailed in the publication and reproduce Figures 1, 3-6:

- go to the code directory, e.g.:

```bash
cd /home/usr/data/acetate_regulation/
```

- open an R session:

```bash
R
```

- run calculations:

```bash
source("run_analysis.R")
```

The code is open-source and available under GPLv3 license.

## Authors
Pierre Millard

## Contact
