## Supplementary file 2 for "Control and regulation of acetate overflow in *Escherichia coli*": run_analysis.R

```
###################
# Set environment #
###################

# load libraries
library(RColorBrewer)
library(gplots)
library(CoRC)

test_khi2 <- function(nb_points, k_val, nb_par){
  # Perform Khi2 statistical test.
  #
  # Args:
  #   nb_points (int): number of data points
  #   k_val (float): khi2 value (cost)
  #   nb_par (int): number of free parameters
  #
  # Returns (list):
  #   $'khi2 value' (float): khi2 value (cost)
  #   $'data points' (int): number of data points
  #   $'fitted parameters' (int): number of free parameters
  #   $'degrees of freedom' (int): degrees of freedom
  #   $'khi2 reduced value' (float): chi2 reduced value
  #   $'p-value, i.e. P(X^2<=value)' (float): p value
  #   $conclusion (str): message indicating whether the models fits (or not) the data at 95% confidence interval
  #
  df <- nb_points - nb_par
  p_val <- pchisq(k_val, df=df)
  khi2test <- list("khi2 value"                  = k_val,
                   "data points"                 = nb_points,
                   "fitted parameters"           = nb_par,
                   "degrees of freedom"          = df,
                   "khi2 reduced value"          = k_val/df,
                   "p-value, i.e. P(X^2<=value)" = p_val)
  if (p_val > 0.95){
    khi2test$conclusion <- "At level of 95% confidence, the model does not fit the data good enough with respect to the provided measurement SD."
  }else{
    khi2test$conclusion <- "At level of 95% confidence, the model fits the data good enough with respect to the provided measurement SD."
  }
  return(khi2test)
}

plot_points <- function(x, y, sd, col="black", offset=1.5, mode="v", cex=1){
  # Scatterplot with error bars.
  #
  # Args:
  #   x (vector): x coordinates
  #   y (vector): y coordinates
  #   sd (vector): error bars
  #   col (color code): color of points
  #   offset (float): width (or height if mode = 'v') of error bars
  #   mode ('v' or 'h'): errors of y (if mode='v') or x (if mode='h')
  #
  if (mode == "v"){
    segments(x0=x, y0=y-sd, x1=x, y1=y+sd) 
    segments(x0=x-offset, y0=y+sd, x1=x+offset, y1=y+sd) 
    segments(x0=x-offset, y0=y-sd, x1=x+offset, y1=y-sd) 
  }else if (mode == "h"){
    segments(x0=x-sd, y0=y, x1=x+sd, y1=y) 
    segments(x0=x+sd, y0=y-offset, x1=x+sd, y1=y+offset) 
    segments(x0=x-sd, y0=y-offset, x1=x-sd, y1=y+offset) 
  }
  points(x, y, pch=21, bg=col, col="black", cex=cex)
}

lines_threshold <- function(x, y, threshold, new, ...){
  # Split data according to a given threshold, and plot lines for
  # each set.
  #
  # Args:
  #   x (vector): x coordinates
  #   y (vector): y coordinates
  #   threshold (float): value of x at which lines should not
  #                      be connected
  #   new (bool): create a new plot if TRUE, otherwise add
  #               lines to an existing plot
  #
  id_1 <- (x < threshold)
  id_2 <- (x > threshold)
  if (new){
    plot(x[id_1], y[id_1], ...)
  }else{
    suppressWarnings(lines(x[id_1], y[id_1], ...))
  }
  suppressWarnings(lines(x[id_2], y[id_2], ...))
}


####################################
# Supplementary information folder #
####################################

# We assume to be already in the working directory, otherwise update the path of the supplementary data folder
#wd <- "C:/Users/millard/Documents/GIT/acetate_regulation/acetate_regulation/"
wd <- getwd()

model_dir <- file.path(wd, "model", "cps")
results_dir <- file.path(wd, "results")


######################
# Model construction #
######################

# Test whether the different models (1-4) fit the data accurately, based on chi2 statistics

# nb_points: number of experimental data points (152)
# nb_par: number of free parameters

# Go to models directory
setwd(model_dir)

# Model 1: no inhibition by acetate
loadModel("Millard2020_Ecoli_glc_ace_model_1.cps")
res_PE_model1 <- runParameterEstimation(method = "Statistics", model = getCurrentModel())
res_PE_model1$chi2 <- test_khi2(nb_points=152, k_val=res_PE_model1$main$objective_value, nb_par=8)

print("chi2-test for model 1:")
print(paste("  p-value = ", res_PE_model1$chi2$`p-value, i.e. P(X^2<=value)`, sep=""))
print(paste("  conclusion: ", res_PE_model1$chi2$conclusion, sep=""))     

# Model 2: inhibition of glycolysis
loadModel("Millard2020_Ecoli_glc_ace_model_2.cps")
res_PE_model2 <- runParameterEstimation(method = "Statistics", model = getCurrentModel())
res_PE_model2$chi2 <- test_khi2(nb_points=152, k_val=res_PE_model2$main$objective_value, nb_par=9)

print("chi2-test for model 2:")
print(paste("  p-value = ", res_PE_model2$chi2$`p-value, i.e. P(X^2<=value)`, sep=""))
print(paste("  conclusion: ", res_PE_model2$chi2$conclusion, sep=""))     

# Model 3: inhibition of TCA cycle
loadModel("Millard2020_Ecoli_glc_ace_model_3.cps")
res_PE_model3 <- runParameterEstimation(method = "Statistics", model = getCurrentModel())
res_PE_model3$chi2 <- test_khi2(nb_points=152, k_val=res_PE_model3$main$objective_value, nb_par=9)

print("chi2-test for model 3:")
print(paste("  p-value = ", res_PE_model3$chi2$`p-value, i.e. P(X^2<=value)`, sep=""))
print(paste("  conclusion: ", res_PE_model3$chi2$conclusion, sep=""))     

# Model 4: inhibition of TCA and glycolysis
loadModel("Millard2020_Ecoli_glc_ace_model_4.cps")
res_PE_model4 <- runParameterEstimation(method = "Statistics", model = getCurrentModel())
res_PE_model4$chi2 <- test_khi2(nb_points=152, k_val=res_PE_model4$main$objective_value, nb_par=10)

print("chi2-test for model 4:")
print(paste("  p-value = ", res_PE_model4$chi2$`p-value, i.e. P(X^2<=value)`, sep=""))
print(paste("  conclusion: ", res_PE_model4$chi2$conclusion, sep=""))     

# plot model evaluation results
setwd(results_dir)
pdf(file="model_construction.pdf", width = 5, height = 5)

barplot(c(res_PE_model1$main$objective_value,
       res_PE_model2$main$objective_value,
       res_PE_model3$main$objective_value,
       res_PE_model4$main$objective_value),
       las=1,
       ylab="variance-weighted SSR",
       names.arg=c("model1", "model2", "model3", "model4"))
chi2_threshold <- 171
abline(h=chi2_threshold)

dev.off()

# plot fitting results for model 4
setwd(model_dir)
experiments <- list("ace_1" = list("conc_ini" = c("ace"=0.92, "glc"=12.9, "X"=0.0671),
                                   "meas" = read.table("data_1mM.txt", header=TRUE, sep="\t")),
                    "ace_10" = list("conc_ini" = c("ace"=9, "glc"=14.192, "X"=0.07),
                                    "meas" = read.table("data_10mM.txt", header=TRUE, sep="\t")),
                    "ace_30" = list("conc_ini" = c("ace"=30.2448, "glc"=12.5, "X"=0.07227),
                                    "meas" = read.table("data_30mM.txt", header=TRUE, sep="\t")))

loadModel("Millard2020_Ecoli_glc_ace_model_4.cps")
setwd(results_dir)
deleteEvent(getEvents()$key)

for (i in c("ace_1", "ace_10", "ace_30")){
  setSpecies(key="Ace0_out", initial_concentration=experiments[[i]]$conc_ini["ace"])
  setSpecies(key="Glc", initial_concentration=experiments[[i]]$conc_ini["glc"])
  setSpecies(key="X", initial_concentration=experiments[[i]]$conc_ini["X"])
  applyInitialState()
  experiments[[i]]$simulations <- runTimeCourse(duration=max(experiments[[i]]$meas$time), suppress_output_before=0.01)$result
}

sd_exp <- c("X"=0.045, "Glc"=0.5, "Ace"=0.2, "Ace_enr"=0.03)

pdf(file = "model_fit.pdf", width = 7, height = 9)
par(mfrow=c(4,3))

plot(experiments$ace_1$simulations$Time, experiments$ace_1$simulations$X, type="l", lwd=2, las=1, xlim=c(0,5), ylim=c(0,1), xaxs="i", yaxs="i", ylab="[biomass] (gDW/L)", xlab="", col="#1F4E79", main="[acetate]=1mM")
plot_points(experiments$ace_1$meas$time, experiments$ace_1$meas$X, sd_exp["X"], offset=0.03, col="#1F4E79", cex=1.2)
plot(experiments$ace_10$simulations$Time, experiments$ace_10$simulations$X, type="l", lwd=2, col="#2E75B6", las=1, xlim=c(0,5), ylim=c(0,1), xaxs="i", yaxs="i", ylab="", xlab="", main="[acetate]=10mM")
plot_points(experiments$ace_10$meas$time, experiments$ace_10$meas$X, sd_exp["X"], offset=0.03, col="#2E75B6", cex=1.2)
plot(experiments$ace_30$simulations$Time, experiments$ace_30$simulations$X, type="l", lwd=2, col="#9DC3E6", las=1, xlim=c(0,5), ylim=c(0,1), xaxs="i", yaxs="i", ylab="", xlab="", main="[acetate]=30mM")
plot_points(experiments$ace_30$meas$time, experiments$ace_30$meas$X, sd_exp["X"], offset=0.03, col="#9DC3E6", cex=1.2)

plot(experiments$ace_1$simulations$Time, experiments$ace_1$simulations$Glc, type="l", lwd=2, las=1, xlim=c(0,5), ylim=c(0,15), xaxs="i", yaxs="i", ylab="[Glc] (mM)", xlab="", col="#548235")
plot_points(experiments$ace_1$meas$time, experiments$ace_1$meas$Glc, sd_exp["Glc"], offset=0.03, col="#548235", cex=1.2)
plot(experiments$ace_10$simulations$Time, experiments$ace_10$simulations$Glc, type="l", lwd=2, col="#70AD47", las=1, xlim=c(0,5), ylim=c(0,15), xaxs="i", yaxs="i", ylab="", xlab="")
plot_points(experiments$ace_10$meas$time, experiments$ace_10$meas$Glc, sd_exp["Glc"], offset=0.03, col="#70AD47", cex=1.2)
plot(experiments$ace_30$simulations$Time, experiments$ace_30$simulations$Glc, type="l", lwd=2, col="#A9D18E", las=1, xlim=c(0,5), ylim=c(0,15), xaxs="i", yaxs="i", ylab="", xlab="")
plot_points(experiments$ace_30$meas$time, experiments$ace_30$meas$Glc, sd_exp["Glc"], offset=0.03, col="#A9D18E", cex=1.2)

plot(experiments$ace_1$simulations$Time, experiments$ace_1$simulations$Acet_out, type="l", lwd=2, las=1, xlim=c(0,5), ylim=c(0,5), xaxs="i", yaxs="i", ylab="[Ace] (mM)", xlab="", col="#B63A2D")
plot_points(experiments$ace_1$meas$time, experiments$ace_1$meas$Ace, sd_exp["Ace"], offset=0.03, col="#B63A2D", cex=1.2)
plot(experiments$ace_10$simulations$Time, experiments$ace_10$simulations$Acet_out, type="l", lwd=2, col="#D6685C", las=1, xlim=c(0,5), ylim=c(8,11), xaxs="i", yaxs="i", ylab="", xlab="")
plot_points(experiments$ace_10$meas$time, experiments$ace_10$meas$Ace, sd_exp["Ace"], offset=0.03, col="#D6685C", cex=1.2)
plot(experiments$ace_30$simulations$Time, experiments$ace_30$simulations$Acet_out, type="l", lwd=2, col="#E49890", las=1, xlim=c(0,5), ylim=c(27,32), xaxs="i", yaxs="i", ylab="", xlab="")
plot_points(experiments$ace_30$meas$time, experiments$ace_30$meas$Ace, sd_exp["Ace"], offset=0.03, col="#E49890", cex=1.2)

plot(experiments$ace_1$simulations$Time, experiments$ace_1$simulations$`Values[Ace_enr]`, type="l", lwd=2, las=1, xlim=c(0,5), ylim=c(0,1), xaxs="i", yaxs="i", ylab="acetate labeling (%)", xlab="time (h)", col="#BF9000")
plot_points(experiments$ace_1$meas$time, experiments$ace_1$meas$Ace_enr, sd_exp["Ace_enr"], offset=0.03, col="#BF9000", cex=1.2)
plot(experiments$ace_10$simulations$Time, experiments$ace_10$simulations$`Values[Ace_enr]`, type="l", lwd=2, col="#FFD966", las=1, xlim=c(0,5), ylim=c(0,0.7), xaxs="i", yaxs="i", ylab="", xlab="time (h)")
plot_points(experiments$ace_10$meas$time, experiments$ace_10$meas$Ace_enr, sd_exp["Ace_enr"], offset=0.03, col="#FFD966", cex=1.2)
plot(experiments$ace_30$simulations$Time, experiments$ace_30$simulations$`Values[Ace_enr]`, type="l", lwd=2, col="#FFE699", las=1, xlim=c(0,5), ylim=c(0,0.3), xaxs="i", yaxs="i", ylab="", xlab="time (h)")
plot_points(experiments$ace_30$meas$time, experiments$ace_30$meas$Ace_enr, sd_exp["Ace_enr"], offset=0.03, col="#FFE699", cex=1.2)

dev.off()


#####################
# Testing the model #
#####################

pdf(file="model_validation.pdf", width = 7, height = 9)
par(mfrow=c(4,3))

# Pulse experiment (Enjalbert et al., 2017)
###########################################

setwd(model_dir)
loadModel("Millard2020_Ecoli_glc_ace_kinetic_model.cps")
setwd(results_dir)

# set biomass concentration at which the pulse is performed
setGlobalQuantities(key = "_X_conc_pulse", initial_value = 0.9)

# simulate response to acetate pulse
res_pulse <- runTimeCourse()

# get simulation results
id_start <- which(res_pulse$result$Ace_out >= 30)[1]+1
id_end <- which(res_pulse$result$Time >= (res_pulse$result$Time[id_start]+8/60))[1]+1

t_pulse <- (res_pulse$result$Time[seq(id_start, id_end)] - res_pulse$result$Time[id_start])
ace_conc_pulse <- res_pulse$result$Ace_out[seq(id_start, id_end)] - res_pulse$result$Ace_out[id_start]
glc_conc_pulse <- res_pulse$result$Glc[seq(id_start, id_end)] - res_pulse$result$Glc[id_start]

# measurements
time_meas <- seq(0,8)/60
glc <- c(0, -0.06931183, -0.151415145, -0.189227994, -0.269451057, -0.290764495, -0.230785281, -0.464084162, -0.551878527)
sd_glc <- c(0, 0.032476344, 0.073133915, 0.113018846, 0.049485284, 0.005325541, 0.163377704, 0.034786419, 0.048477157)
ace <- c(0, -0.027907926, -0.078000853, -0.155334163, -0.165031608, -0.111098424, -0.182877548, -0.237262298, -0.276903255)
sd_ace <- c(0, 0.002740145, 0.025693594, 0.053641876, 0.089975321, 0.005875669, 0.085604161, 0.061930626, 0.099140975)

# plot simulations vs measurements
plot(t_pulse, ace_conc_pulse, type="l", ylim=c(-0.4,0), las=1, main="Enjalbert_2017 (pulse 30mM ace)", ylab="change of [Ace]", xlab="time (h)", col="#D6685C", lwd=2)
plot_points(time_meas, ace, sd_ace, offset=0.002, col="#D6685C")
plot(t_pulse, glc_conc_pulse, type="l", main="Enjalbert_2017 (pulse 30mM ace)", las=1, ylab="change of [Glc]", xlab="time (h)", col="#70AD47", lwd=2)
plot_points(time_meas, glc, sd_glc, offset=0.002, col="#70AD47")

# simulate control experiment (i.e. no acetate pulse)
deleteEvent(getEvents()$key)

res_nopulse <- runTimeCourse()

# get simulation results
ace_conc_nopulse <- res_nopulse$result$Ace_out[seq(id_start, id_end)] - res_nopulse$result$Ace_out[id_start]
glc_conc_nopulse <- res_nopulse$result$Glc[seq(id_start, id_end)] - res_nopulse$result$Glc[id_start]

# measurements
glc_nopulse <- c(0, -0.195637774, -0.325512845, -0.433785334, -0.628518958, -0.726913548, -0.892908748, -1.071230579, -1.16285575)
sd_glc_nopulse <- c(0, 0.058023617, 0.109115511, 0.047090371, 0.052331837, 0.065619906, 0.133896355, 0.16828754, 0.03515465)
ace_nopulse <- c(0, 0.01012067, 0.05974009, 0.086787283, 0.086690257, 0.104329693, 0.128507087, 0.130026354, 0.187336965)
sd_ace_nopulse <- c(0, 0.005117257, 0.022507218, 0.040319045, 0.037548873, 0.037235285, 0.044494365, 0.045029023, 0.023982374)

# plot simulations vs measurements
plot(t_pulse, ace_conc_nopulse, type="l", ylim=c(0,0.3), main="Enjalbert_2017 (control)", las=1, ylab="change of [Ace]", xlab="time (h)", col="#D6685C", lwd=2)
plot_points(time_meas, ace_nopulse, sd_ace_nopulse, offset=0.002, col="#D6685C")
plot(t_pulse, glc_conc_nopulse, type="l", ylim=c(-1.4,0), main="Enjalbert_2017 (control)", las=1, ylab="change of [Glc]", xlab="time (h)", col="#70AD47", lwd=2)
plot_points(time_meas, glc_nopulse, sd_glc_nopulse, offset=0.002, col="#70AD47")


# Chemostat experiment (Renilla et al., 2012)
#############################################

setwd(model_dir)
loadModel("Millard2020_Ecoli_glc_ace_kinetic_model.cps")
setwd(results_dir)

# delete events and set glc feed to 1
deleteEvent(getEvents()$key)
setGlobalQuantities(key = "_feed", initial_value = 1)


n_step <- 50
dilution_rates <- seq(0.1, 0.5, length.out = n_step)
fluxes <- c("Values[v_growth_rate]", "Values[v_glc_uptake]", "Values[v_ace_net]")
res_chemostat <- matrix(NA, nrow=n_step, ncol=length(fluxes)+1, dimnames=list(r=NULL, c=c("dilution_rate", fluxes)))
for (i in seq(n_step)){
  setGlobalQuantities(key = "_dilution_rate", initial_value = dilution_rates[i])
  res_ss <- runSteadyState()
  res_chemostat[i,] <- c(dilution_rates[i], unlist(res_ss$global_quantities[res_ss$global_quantities$key %in% fluxes, "value"]))
}

# acetate flux as function of dilution rate
dilution_rate <- c(0.09586056644880175, 0.20043572984749464, 0.2997821350762528, 0.3468409586056645, 0.39912854030501094, 0.44618736383442276, 0.5002178649237474)
q_ace <- c(0.40472342596168076, 0.7396156614846294, 1.477019302736899, 1.2154669794005626, 1.96934635755591, 2.0929143843289824, 2.006569318707304)
q_ace_sd <- c(0.120440687823878, 0.15316333195082, 0.04595662724122, 0.0274214232252601, 0.28885933172199, 0.0777639526513201, 0.28664731148963)

plot(res_chemostat[,"dilution_rate"], res_chemostat[,"Values[v_ace_net]"], type="l", las=1, xlim=c(0,0.6), ylim=c(0,4), main="Renilla_2012", lwd=2, xlab="dilution rate (h-1)", ylab="ace flux (mmol/gDW/h)", col="#D6685C")
plot_points(dilution_rate, q_ace, q_ace_sd, offset=0.01, col="#D6685C")

# acetate flux as function of glc uptake
q_glc <- c(1.43478260869565, 2.7391304347826, 4.59130434782608, 4.69565217391304, 5.7391304347826, 5.92173913043478, 6.10434782608695, 7.12173913043477, 8.34782608695652)
q_ace <- c(0.381358340437624, 0.762906128635029, 1.40509614473808, 1.17836506583309, 1.99602159704461, 2.08290233967983, 2.01292033721701, 2.44717249218528, 2.5848252344416)
q_ace_sd <- c(0.139547219854126, 0.139471440750213, 0.15697641375391, 0.1045751633987, 0.19179691200152, 0.0174291938997801, 0.24397082504499, 0.52295159609737, 0.33130624230368)
q_glc_sd <- c(0.20234760540444, 0.58899555210581, 0.36827270271402, 0.14607356337583, 0.630432119199, 0.13069807968161, 0.47630364392495, 0.99361847036576, 0.66185122193432)

plot(res_chemostat[,"Values[v_glc_uptake]"], res_chemostat[,"Values[v_ace_net]"], type="l", las=1, xlim=c(0,10), ylim=c(0,4), main="Renilla_2012", lwd=2, xlab="glc uptake (mmol/gDW/h)", ylab="ace flux (mmol/gDW/h)", col="#D6685C")
plot_points(q_glc, q_ace, q_ace_sd, offset=0.2, col="#D6685C")
plot_points(q_glc, q_ace, q_glc_sd, offset=0.08, mode="h", col="#D6685C")


# Steady-state fluxes for acetate concentration between 0.1 and 100mM (Enjalbert et al., 2017; Pinhal et al., 2019)
###################################################################################################################

setwd(model_dir)
loadModel("Millard2020_Ecoli_glc_ace_kinetic_model.cps")
setwd(results_dir)

# remove events and fix concentrations of actate, glucose and biomass
deleteEvent(getEvents()$key)
setSpecies(key="Ace_out", type="fixed")
setSpecies(key="Glc", type="fixed")
setSpecies(key="X", type="fixed")

# run simulations
n_step <- 50
ace_concentration <- 10**seq(-1, 2, length.out = n_step)
fluxes <- c("Values[v_growth_rate]", "Values[v_glc_uptake]", "Values[v_ace_net]")
res_ace_range <- matrix(NA, nrow=n_step, ncol=length(fluxes)+1, dimnames=list(r=NULL, c=c("ace_concentration", fluxes)))
for (i in seq(n_step)){
  setSpecies(key="Ace_out", initial_concentration=ace_concentration[i])
  applyInitialState()
  res_ss <- runSteadyState()
  res_ace_range[i,] <- c(ace_concentration[i], unlist(res_ss$global_quantities[res_ss$global_quantities$key %in% fluxes, "value"]))
}

xlab_main <- c(0.1, 1, 10, 100)
xlab_sec <- c(seq(0.2, 0.9, by=0.1), seq(2, 9, by=1), seq(20, 90, by=10))

# growth rate as function of acetate concentration
growth_rates <- c(0.521128511, 0.611148842625582, 0.613161998174498, 0.502533817, 0.496290415, 0.488201506, 0.547635665, 0.499830448, 0.474554197, 0.425356578, 0.377534684, 0.645724326, 0.618475601, 0.554887936, 0.564811523, 0.527571192, 0.434972836, 0.3824734, 0.583623355, 0.620905534, 0.564259247, 0.532148135, 0.483885215, 0.557074418, 0.630654409249223)
sd_growth_rates <- c(0.001793104, 0.00204807928657914, 0.00219396182484705, 0.001709207, 0.001846205, 0.001757403, 0.001821375, 0.001025702, 0.001940912, 0.001204707, 0.001999188, 0.001418374, 0.001932601, 0.001455791, 0.001574234, 0.001206265, 0.001292476, 0.001068259, 0.001804648, 0.001643459, 0.001598405, 0.001121218, 0.000912408, 0.00194896, 0.00203369597574686)
ace_conc <- c(0.451964624, 1.11600286648471, 2.04718732931708, 2.86252145, 5.285907977, 9.106164204, 16.67476528, 16.76626787, 17.00218707, 30.8667961, 57.92292091, 0.478352574, 4.55673229, 8.10163028, 8.22100734, 7.829591756, 33.53244905, 66.20361403, 0.436956014, 1.28468189, 1.555875222, 12.02564968, 30.24566673, 0.548011282, 2.29562227069566)
sd_ace_conc <- c(0.262318006, 0.281361538208953, 0.289527601555302, 0.302351163, 0.327705513, 0.330782201, 0.277259011, 0.233956798, 0.31883929, 0.300578057, 0.313371784, 0.231202155, 0.264687437, 0.243480317, 0.289821733, 0.263808862, 0.289478134, 0.264861034, 0.22461248, 0.229031308, 0.241718918, 0.254111384, 0.187394292, 0.011107606, 0.290519090995892)
plot(res_ace_range[,"ace_concentration"], xaxt='n', res_ace_range[,"Values[v_growth_rate]"], las=1, col="#2E75B6", lwd=2, type="l", xlim=c(0.1,100), ylim=c(0,0.8), log="x", main="Enjalbert_2017", xlab="[acetate] (mM)", ylab="growth rate (h-1)")
axis(side = 1, at = xlab_main, labels = TRUE)
axis(side = 1, at = xlab_sec, labels = FALSE, tcl=-0.3)
plot_points(ace_conc, growth_rates, sd_growth_rates, offset=0.01, col="#2E75B6")
plot_points(ace_conc, growth_rates, sd_ace_conc, offset=0.02, mode="h", col="#2E75B6")

# glc uptake as function of acetate concentration
glc_upt <- c(8.654860167, 8.36127542981722, 7.98010111285252, 9.236935826, 8.274418986, 7.560431219, 7.339194455, 5.775312502, 6.423391263, 5.1544758, 3.938631573, 8.115447647, 9.28067031, 6.737153424, 7.172748804, 5.884186033, 5.684201497, 4.811576974, 9.632702365, 8.055042777, 9.708342814, 7.100081588, 5.505759496, 9.242859752, 8.18621623190759)
sd_glc_upt <- c(0.337812425, 0.38531328268303, 0.373770045721031, 0.356787032, 0.334672954, 0.317509322, 0.288025925, 0.16053276, 0.375934255, 0.293148172, 0.359225607, 0.197331684, 0.360984112, 0.229372278, 0.241820396, 0.20450532, 0.260869273, 0.216134352, 0.34289286, 0.350305744, 0.293144783, 0.220135755, 0.153471508, 0.25245346, 0.396815184905029)
ace_conc <- c(0.451964624, 1.11600286648471, 2.04718732931708, 2.86252145, 5.285907977, 9.106164204, 16.67476528, 16.76626787, 17.00218707, 30.8667961, 57.92292091, 0.478352574, 4.55673229, 8.10163028, 8.22100734, 7.829591756, 33.53244905, 66.20361403, 0.436956014, 1.28468189, 1.555875222, 12.02564968, 30.24566673, 0.548011282, 2.29562227069566)
sd_ace_conc <- c(0.262318006, 0.281361538208953, 0.289527601555302, 0.302351163, 0.327705513, 0.330782201, 0.277259011, 0.233956798, 0.31883929, 0.300578057, 0.313371784, 0.231202155, 0.264687437, 0.243480317, 0.289821733, 0.263808862, 0.289478134, 0.264861034, 0.22461248, 0.229031308, 0.241718918, 0.254111384, 0.187394292, 0.011107606, 0.290519090995892)
plot(res_ace_range[,"ace_concentration"], res_ace_range[,"Values[v_glc_uptake]"], xaxt='n', las=1, type="l", lwd=2, xlim=c(0.1,100), col="#70AD47", ylim=c(0,10), log="x", main="Enjalbert_2017", xlab="[acetate] (mM)", ylab="glc uptake (mmol/gDW/h)")
axis(side = 1, at = xlab_main, labels = TRUE)
axis(side = 1, at = xlab_sec, labels = FALSE, tcl=-0.3)
plot_points(ace_conc, glc_upt, sd_glc_upt, offset=0.02, col="#70AD47")
plot_points(ace_conc, glc_upt, sd_ace_conc, offset=0.2, mode="h", col="#70AD47")

# ace flux as function of acetate concentration
ace_flx <- c(3.5, -2.7, 1.516999356, 1.26845123082679, 0.775821380507016, 0.678877137, 0.017366464, -0.991478151, -1.286687213, -2.078474994, -1.530841439, -1.525342269, -1.253581266, 1.984679487, 0.546462624, -0.136780389, -0.393883917, -0.610240984, -1.120767885, -1.277455315, 2.574285211, 2.051935093, 1.828415596, -1.262442483, -1.317987733, 2.333568565, 1.85234639824858)
sd_ace_flx <- c(0.35, 0.27, 0.316118066, 0.388752117258161, 0.40715278851436, 0.33357638, 0.37333751, 0.347029894, 0.280501612, 0.195031303, 0.376252463, 0.226182385, 0.303661317, 0.253610517, 0.385450715, 0.243880325, 0.30665695, 0.257983739, 0.23844407, 0.198458448, 0.299832036, 0.334956504, 0.317134334, 0.263807154, 0.195219648, 0.016120887, 0.386174129654754)
ace_conc <- c(0.2, 100, 0.451964624, 1.11600286648471, 2.04718732931708, 2.86252145, 5.285907977, 9.106164204, 16.67476528, 16.76626787, 17.00218707, 30.8667961, 57.92292091, 0.478352574, 4.55673229, 8.10163028, 8.22100734, 7.829591756, 33.53244905, 66.20361403, 0.436956014, 1.28468189, 1.555875222, 12.02564968, 30.24566673, 0.548011282, 2.29562227069566)
sd_ace_conc <- c(0.05, 10, 0.262318006, 0.281361538208953, 0.289527601555302, 0.302351163, 0.327705513, 0.330782201, 0.277259011, 0.233956798, 0.31883929, 0.300578057, 0.313371784, 0.231202155, 0.264687437, 0.243480317, 0.289821733, 0.263808862, 0.289478134, 0.264861034, 0.22461248, 0.229031308, 0.241718918, 0.254111384, 0.187394292, 0.011107606, 0.290519090995892)
plot(res_ace_range[,"ace_concentration"], res_ace_range[,"Values[v_ace_net]"], xaxt='n', las=1, type="l", lwd=2, xlim=c(0.1,100), ylim=c(-4,4), log="x", main="Enjalbert_2017, Pinhal_2019", col="#D6685C", xlab="[acetate] (mM)", ylab="ace flux (mmol/gDW/h)")
abline(h=0)
axis(side = 1, at = xlab_main, labels = TRUE)
axis(side = 1, at = xlab_sec, labels = FALSE, tcl=-0.3)
plot_points(ace_conc, ace_flx, sd_ace_flx, offset=0.04, col="#D6685C")
plot_points(ace_conc, ace_flx, sd_ace_conc, offset=0.2, mode="h", col="#D6685C")

dev.off()


##############################
# Metabolic control analyses #
##############################

setwd(model_dir)
loadModel("Millard2020_Ecoli_glc_ace_kinetic_model.cps")
setwd(results_dir)

# delete events and fix concentrations of biomass and extracellular glc and acetate
deleteEvent(getEvents()$key)
setSpecies(key="Ace_out", type="fixed")
setSpecies(key="Glc", type="fixed")
setSpecies(key="X", type="fixed")

# calculate flux control coefficients
n_step <- 50
ace_range <- 10**(seq(-1,2,length.out = n_step))
res_ace_MCA <- matrix(NA, nrow=length(ace_range), ncol=7)
res_ace_MCA[,1] <- ace_range
for (i in seq(n_step)){
  setSpecies(key="Ace_out{cell}", initial_concentration = ace_range[i], model = getCurrentModel())
  applyInitialState(model = getCurrentModel())
  res_MCA <- runMetabolicControlAnalysis(model = getCurrentModel())
  if (res_MCA$result_ss != "found"){
    print("error")
  }else{
    fcc <- res_MCA$flux_control_coefficients_scaled
    res_ace_MCA[i,seq(2,7)] <- fcc["(ackA)", colnames(fcc) != "'Summation Error'"]
  }
}
colnames(res_ace_MCA) <- c("ace_conc", colnames(fcc)[colnames(fcc) != "'Summation Error'"])

# plot results
pdf(file = "results_control_1.pdf", height = 4, width = 5)

setSpecies(key="Ace_out{cell}", initial_concentration = 0.1, model = getCurrentModel())
applyInitialState(model = getCurrentModel())
res_MCA <- runMetabolicControlAnalysis(model = getCurrentModel())

hmc <- res_MCA$flux_control_coefficients_scaled[c("(glc_upt)", "(ackA)", "(sink)"), c("(glc_upt)", "(sink)", "(pta)", "(ackA)", "(ace_xch)")]
colnames(hmc) <- c("glycolysis", "TCA", "Pta", "AckA", "Ace_xch")
rownames(hmc) <- c("glycolysis", "acetate", "TCA")
rgb.palette <- colorRampPalette(brewer.pal(n = 11, name = "RdBu"))
breaks <- c(-1, -0.5, -0.2, -0.1, -0.05, -0.01, 0.01, 0.05, 0.1, 0.2, 0.5, 1, 1.5)
col_scale <- rgb.palette(length(breaks)-1)
col_scale[6] <- "#DDDDDD"
heatmap.2(hmc, Rowv=FALSE, Colv=FALSE, dendrogram="none", col=col_scale, breaks=breaks,
          scale="none", trace="none", colsep=seq(0,ncol(hmc)), rowsep=seq(0,nrow(hmc)), sepwidth=c(0.04,0.02))

dev.off()

pdf(file="results_control_2.pdf", width = 7, height = 9)
par(mfrow=c(4,3))

xlab_main <- c(0.1, 1, 10, 100)
xlab_sec <- c(seq(0.2, 0.9, by=0.1), seq(2, 9, by=1), seq(20, 90, by=10))
conc_threshold <- 14.27174

ctrl_by_acetate_pathway <- apply(res_ace_MCA[,c("(ackA)", "(pta)", "(ace_xch)")], 1, sum)
ctrl_by_glc_upt <- res_ace_MCA[,"(glc_upt)"]
ctrl_by_TCA <- res_ace_MCA[,"(sink)"]

plot(x=ace_range, y=ctrl_by_acetate_pathway, type="l", xaxt="n", las=1, ylim=c(0,1), log="x", xaxs="i", yaxs="i", xlab="[acetate] (mM)", ylab="control by acetate pathway", lwd=2)
axis(side = 1, at = xlab_main, labels = TRUE)
axis(side = 1, at = xlab_sec, labels = FALSE, tcl=-0.3)

lines_threshold(x=ace_range, y=ctrl_by_glc_upt, threshold=conc_threshold, new=TRUE, xaxt="n", xaxs="i", yaxs="i", las=1, xlim=c(0.1,100), type="l", ylim=c(-3,3), log="x", xlab="[acetate] (mM)", ylab="control by glycolysis", lwd=2)
axis(side = 1, at = xlab_main, labels = TRUE)
axis(side = 1, at = xlab_sec, labels = FALSE, tcl=-0.3)
abline(h=0)

lines_threshold(x=ace_range, y=ctrl_by_TCA, threshold=conc_threshold, new=TRUE, xaxt="n", xaxs="i", yaxs="i", las=1, xlim=c(0.1,100), type="l", ylim=c(-3,3), log="x", xlab="[acetate] (mM)", ylab="control by TCA", lwd=2)
axis(side = 1, at = xlab_main, labels = TRUE)
axis(side = 1, at = xlab_sec, labels = FALSE, tcl=-0.3)
abline(h=0)

plot(x=ace_range, y=ctrl_by_TCA+ctrl_by_glc_upt, type="l", ylim=c(0,1), xaxt="n", xaxs="i", yaxs="i", las=1, log="x", xlab="[acetate] (mM)", ylab="control by TCA+glycolysis", lwd=2)
axis(side = 1, at = xlab_main, labels = TRUE)
axis(side = 1, at = xlab_sec, labels = FALSE, tcl=-0.3)

plot(x=ace_range, y=-ctrl_by_TCA/ctrl_by_glc_upt, type="l", ylim=c(0,2), xaxt="n", xaxs="i", yaxs="i", las=1, log="x", xlab="[acetate] (mM)", ylab="control by TCA/glycolysis", lwd=2)
axis(side = 1, at = xlab_main, labels = TRUE)
axis(side = 1, at = xlab_sec, labels = FALSE, tcl=-0.3)
abline(h=1)

dev.off()


#################################
# Metabolic regulation analyses #
#################################

n_step <- 300
delta_p <- 0.001
conc_threshold <- 14.27174
ace_range <- 10**(seq(-1, 2, length.out = n_step))
ace_range <- ace_range[abs(ace_range - conc_threshold) > 0.1]
res_ace_regulation <- matrix(NA, nrow=length(ace_range), ncol=4, dimnames=list(r=NULL, c=c("ace_conc", "via_acetate_pathway", "via_glc_upt", "via_tca")))

for (i in seq(length(ace_range))){
  
  # set ace concentration
  setSpecies(key="Ace_out{cell}", initial_concentration = ace_range[i])
  applyInitialState()
  
  # get steady-state
  res_ss_i <- runSteadyState(update_model=TRUE)$global_quantities
  
  # calculate control coefficients
  res_MCA_R <- runMCA()$flux_control_coefficients_scaled
  
  # calculate elasticities
  
  # fix acetylCoA concentration to calculate elasticity of each pathway wrt acetate
  setSpecies(key="AcCoA", type="fixed")
  
  # change acetate concentration
  setSpecies(key="Ace_out{cell}", initial_concentration = ace_range[i]*(1+delta_p))
  applyInitialState()
  
  # get steady-state
  res_ss_i_eps <- runSteadyState()$global_quantities
  
  # calculate elasticities (using the more stable numerical method, both being equivalent)
  #elasticities <- (log(abs(res_ss_i_eps$value)) - log(abs(res_ss_i$value))) / log(1+delta_p)
  #print(elasticities)
  elasticities <- (res_ss_i_eps$value - res_ss_i$value) / delta_p / res_ss_i_eps$value
  #print(elasticities)
  
  # reset balance on accoa
  setSpecies(key="AcCoA", type="reactions")
  
  # calculate response coefficient
  # acetate via Pta-AckA
  res_reg_ace_ace <- sum(res_MCA_R["(ackA)", c("(ackA)", "(pta)", "(ace_xch)")]) * elasticities[res_ss_i_eps$key == "Values[v_ace_net]"]
  # acetate via glc uptake
  res_reg_ace_glc <- res_MCA_R["(ackA)", "(glc_upt)"] * elasticities[res_ss_i_eps$key == "Values[v_glc_uptake]"]
  # acetate via sink
  res_reg_ace_tca <- res_MCA_R["(ackA)", "(sink)"] * elasticities[res_ss_i_eps$key == "Values[v_growth_rate]"]
  
  res_ace_regulation[i,] <- c(ace_range[i], res_reg_ace_ace, res_reg_ace_glc, res_reg_ace_tca)
}


pdf(file="results_regulation.pdf", width = 7, height = 9)
par(mfrow=c(4,3))

xlab_main <- c(0.1, 1, 10, 100)
xlab_sec <- c(seq(0.2, 0.9, by=0.1), seq(2, 9, by=1), seq(20, 90, by=10))
conc_threshold <- 14.27174

# plot partitioned response coefficients
lines_threshold(ace_range, res_ace_regulation[,"via_acetate_pathway"], threshold=conc_threshold, new=TRUE, xaxt="n", las=1, xaxs="i", yaxs="i", col="#2E75B6", xlim=c(0.1,100), type="l", log="x", ylim=c(-5, 5), xlab="[acetate] (mM)", ylab="R_ace_pathway", lwd=2)
axis(side = 1, at = xlab_main, labels = TRUE)
axis(side = 1, at = xlab_sec, labels = FALSE, tcl=-0.3)
abline(h=0)
lines_threshold(ace_range, res_ace_regulation[,"via_glc_upt"], threshold=conc_threshold, new=TRUE, xaxt="n", las=1, xaxs="i", yaxs="i", col="#D6685C", xlim=c(0.1,100), type="l", log="x", ylim=c(-5, 5), xlab="[acetate] (mM)", ylab="R_Glc_uptake", lwd=2)
axis(side = 1, at = xlab_main, labels = TRUE)
axis(side = 1, at = xlab_sec, labels = FALSE, tcl=-0.3)
abline(h=0)
lines_threshold(ace_range, res_ace_regulation[,"via_tca"], threshold=conc_threshold, new=TRUE, xaxt="n", las=1, xaxs="i", yaxs="i", col="#70AD47", xlim=c(0.1,100), type="l", log="x", ylim=c(-5, 5), xlab="[acetate] (mM)", ylab="R_TCA", lwd=2)
axis(side = 1, at = xlab_main, labels = TRUE)
axis(side = 1, at = xlab_sec, labels = FALSE, tcl=-0.3)
abline(h=0)

# plot contribution of each pathway
contribution <- apply(res_ace_regulation[,2:4], 1, FUN=function(x) sum(abs(x)))
lines_threshold(ace_range, res_ace_regulation[,"via_acetate_pathway"]/contribution, threshold=conc_threshold, new=TRUE, xaxt="n", las=1, xaxs="i", yaxs="i", xlim=c(0.1,100), type="l", log="x", xlab="[acetate] (mM)", ylab="relative_R", ylim=c(-0.7, 0.7), lwd=2, col="#2E75B6")
axis(side = 1, at = xlab_main, labels = TRUE)
axis(side = 1, at = xlab_sec, labels = FALSE, tcl=-0.3)
abline(h=0)
lines_threshold(ace_range, res_ace_regulation[,"via_glc_upt"]/contribution, threshold=conc_threshold, new=FALSE, type="l", col="#D6685C", lwd=2)
lines_threshold(ace_range, res_ace_regulation[,"via_tca"]/contribution, threshold=conc_threshold, new=FALSE, type="l", col="#70AD47", lwd=2)

dev.off()
```
