## Supplementary figures and images for "Control and regulation of acetate overflow in *Escherichia coli*"

### model_construction.pdf

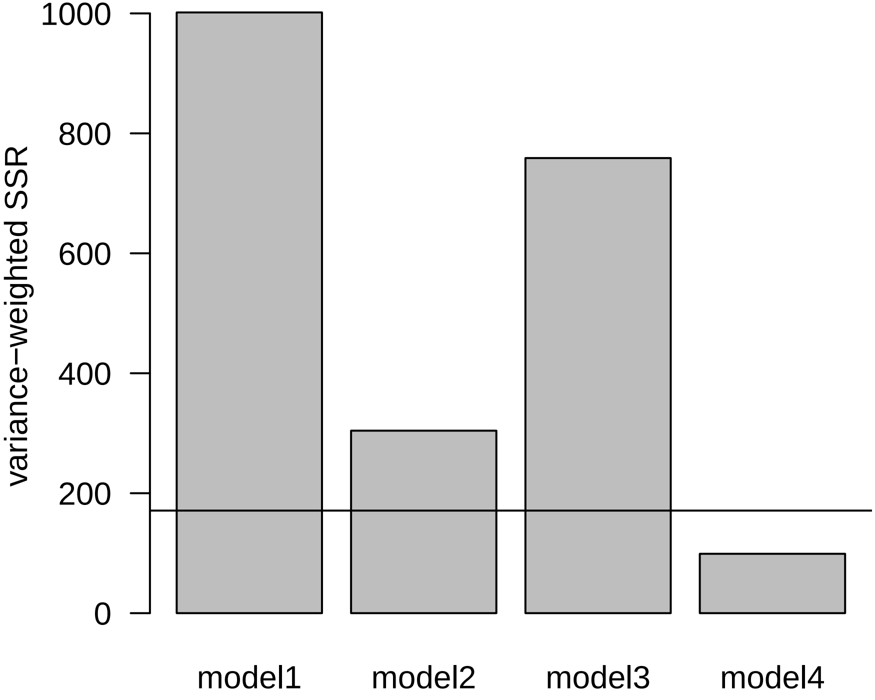

### model_fit.pdf

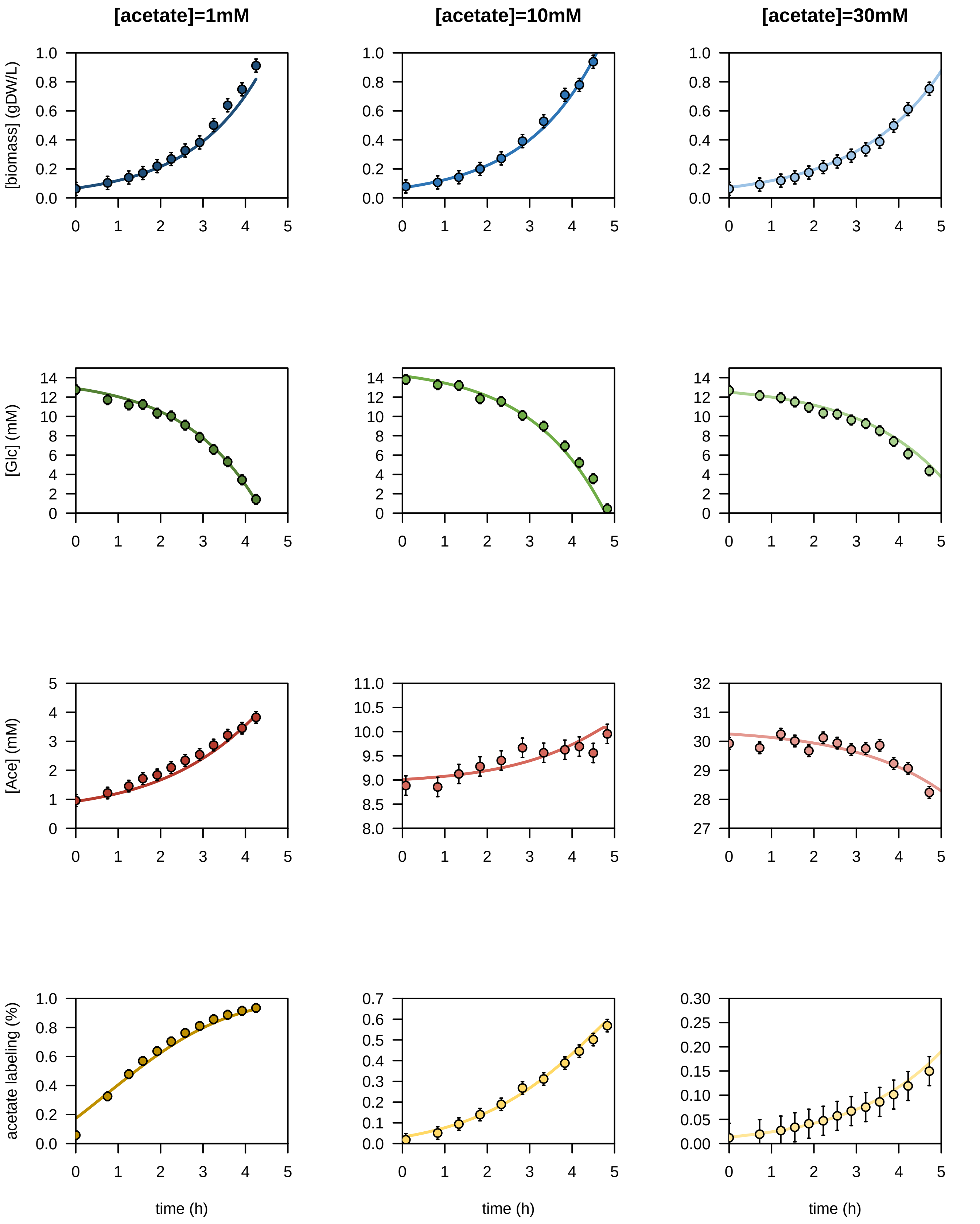

### model_validation.pdf

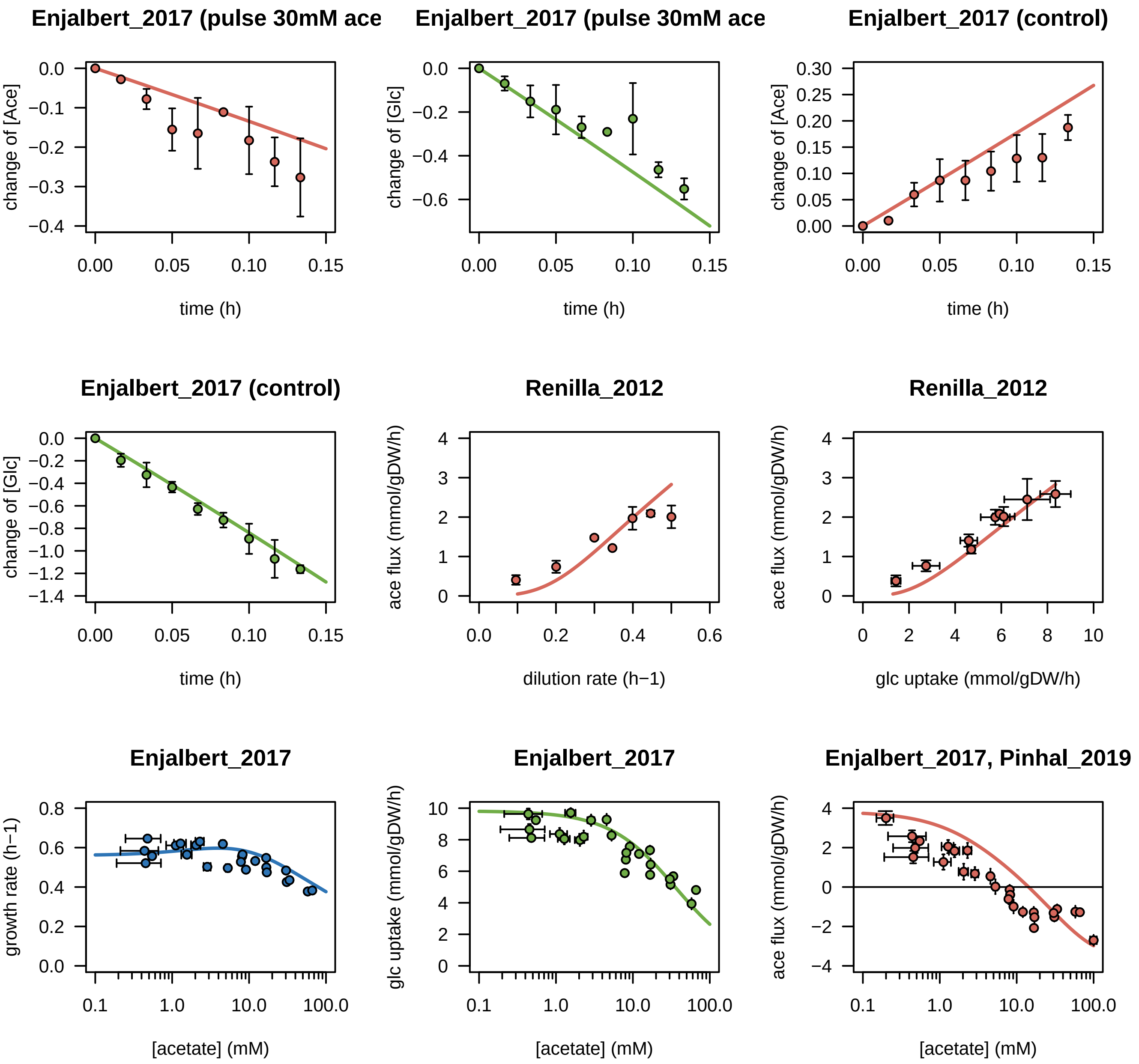

### results_control_1.pdf

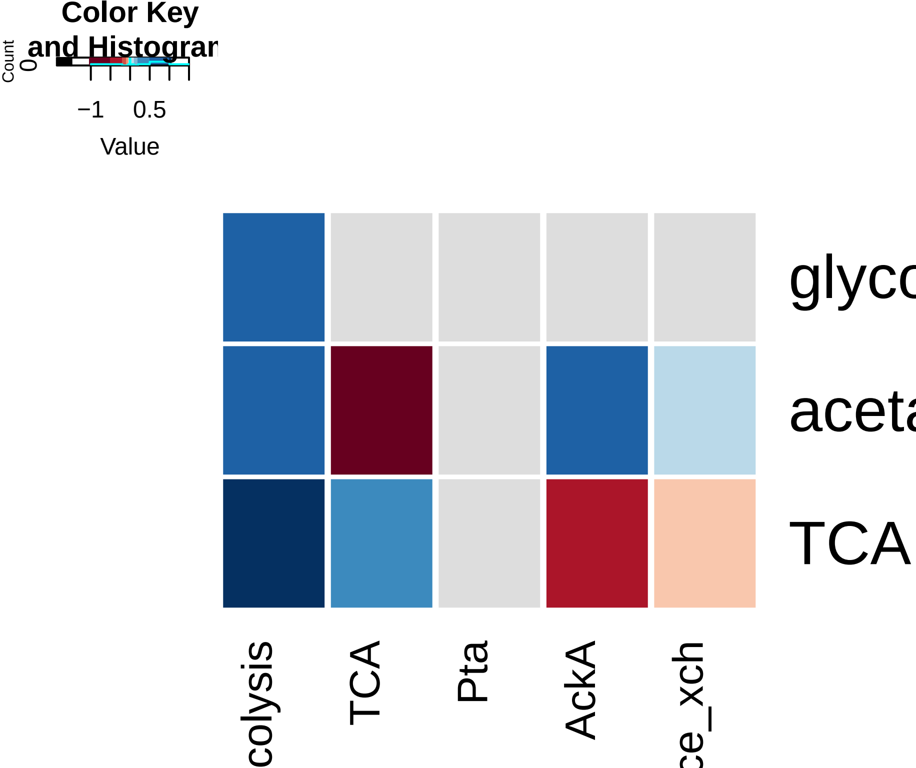

### results_control_2.pdf

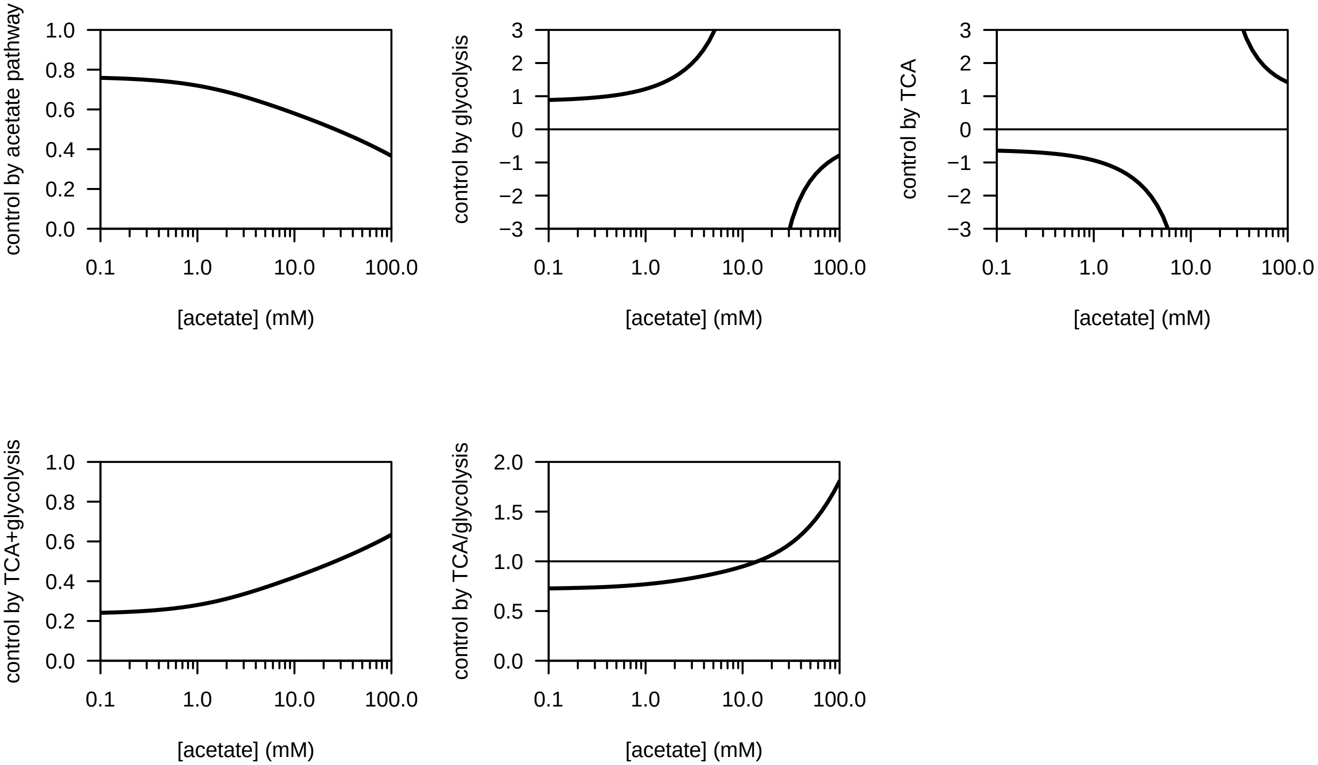

### results_regulation.pdf

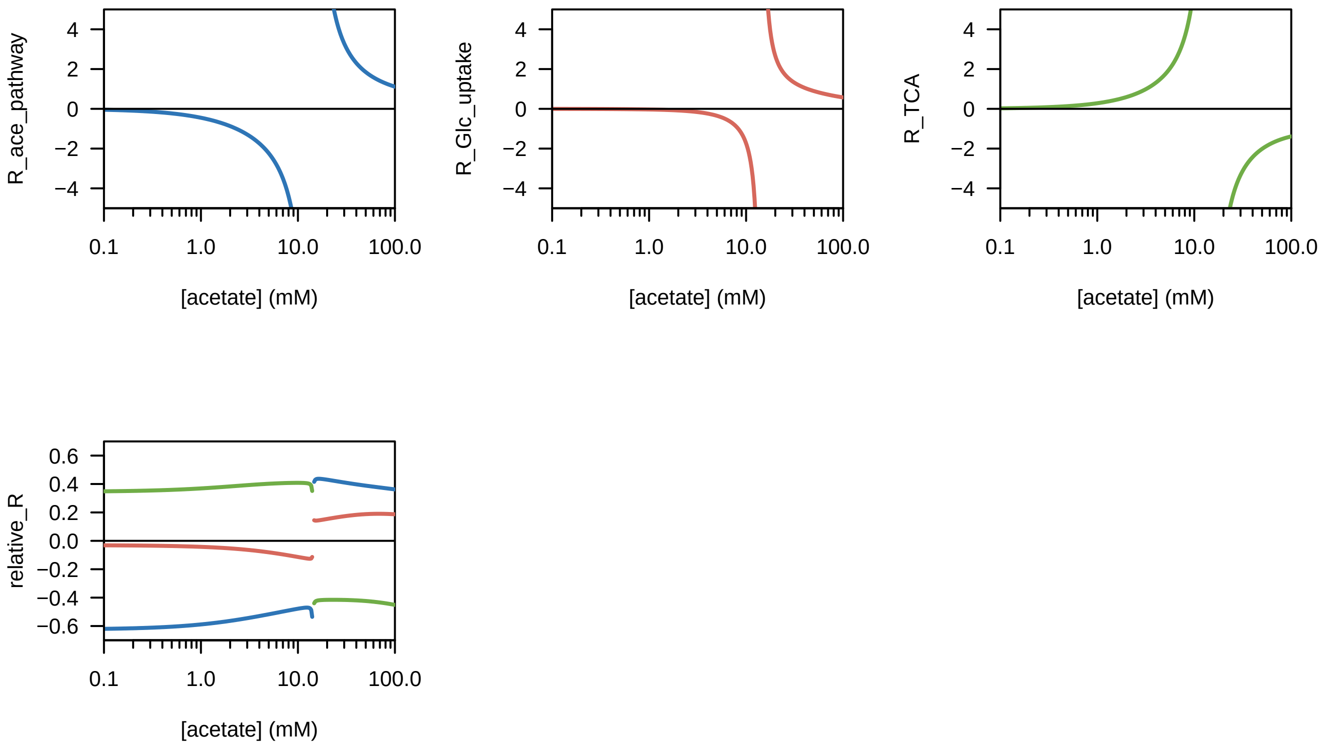
